# Proximity labeling of the Tau repeat domain enriches RNA-binding proteins that are altered in Alzheimer’s disease and related tauopathies

**DOI:** 10.1101/2025.01.22.633945

**Authors:** Sarah M. Shapley, Anantharaman Shantaraman, Masin A. Kearney, Eric B. Dammer, Duc M. Duong, Christine A. Bowen, Pritha Bagchi, Qi Guo, Srikant Rangaraju, Nicholas T. Seyfried

## Abstract

In Alzheimer’s disease (AD) and other tauopathies, tau dissociates from microtubules and forms toxic aggregates that contribute to neurodegeneration. Although some of the pathological interactions of tau have been identified from postmortem brain tissue, these studies are limited by their inability to capture transient interactions. To investigate the interactome of aggregate-prone fragments of tau, we applied an *in vitro* proximity labeling technique using split TurboID biotin ligase (sTurbo) fused with the tau microtubule repeat domain (TauRD), a core region implicated in tau aggregation. We characterized sTurbo TauRD co-expression, robust enzyme activity and nuclear and cytoplasmic localization in a human cell line. Following enrichment of biotinylated proteins and mass spectrometry, we identified over 700 TauRD interactors. Gene ontology analysis of enriched TauRD interactors highlighted processes often dysregulated in tauopathies, including spliceosome complexes, RNA-binding proteins (RBPs), and nuclear speckles. The disease relevance of these interactors was supported by integrating recombinant TauRD interactome data with human AD tau interactome datasets and protein co-expression networks from individuals with AD and related tauopathies. This revealed an overlap with the TauRD interactome and several modules enriched with RBPs and increased in AD and Progressive Supranuclear Palsy (PSP). These findings emphasize the importance of nuclear pathways in tau pathology, such as RNA splicing and nuclear-cytoplasmic transport and establish the sTurbo TauRD system as a valuable tool for exploring the tau interactome.

## Introduction

Tau protein, which is encoded by the microtubule-associated protein tau (*MAPT*) gene^1, 2^, plays a crucial role in maintaining the dynamic nature of the cytoskeleton^3, 4^. The misfolding and accumulation of Tau in the brain is the key characteristic of a group of disorders classified as Tauopathies^5, 6^. In rare instances, mutations in the *MAPT* gene can lead to specific forms of frontotemporal dementia (FTD)^7, 8^. The aggregation of tau can also result in cognitive, behavioral, linguistic, and motor impairments, depending on the specific area of pathology. A significant feature of these conditions, including Alzheimer’s disease (AD) and other tauopathies, is the formation of neurofibrillary tangles (NFTs)^9, 10^, which are strongly correlated to the onset of cognitive decline^11^.

Gain or loss of interaction partners is also an important consequence of disease pathophysiology and may result in ‘rewiring’ of protein-protein interaction (PPI) networks, with associated alteration in function^12^. To this end, in a pathological state, tau dissociates from microtubules and forms NFTs, leading to a loss and aberrant gain of tau PPIs. To identify tau interacting or co-aggregating partners, techniques such as affinity purification (AP) using tau antibodies, along with laser capture microdissection (LCM) and mass spectrometry (MS), have been employed on postmortem brain tissue from AD cases and revealed numerous pathways and interactors associated with tau, highlighting its roles in translation, energy metabolism, long-term potentiation, and mitochondrial function^13–16^. Moreover, within large-scale proteomic analysis of human post-mortem brain tissues, RBPs have emerged as a class of proteins increased in abundance in AD compared to controls and enriched in modules that correlate with tau tangle pathology and splicing defects^17–19^. To this end, several lines of evidence have demonstrated that tau interacts with RBPs under both normal and pathological conditions^20, 21^. For example, tau immunoprecipitated (IP) from AD brain interacts with RBPs, including U1 snRNP proteins (U1-70K, U1A)^16^. Many of these core RBPs, including U1-70K, mis-localize to the neuronal cytoplasm and co-aggregate with AD tau^22, 23^. These findings have been confirmed through various mass spectrometry methods and validated by immunohistochemistry in human AD brain^24, 25^. Other RBPs found in tau aggregates include SRRM2^26^, PNN^24^, TIA1^27–30^, HNRNPA2B1^31^, Musashi 1/2^32^, SFPQ^33^, and serine/arginine-rich splicing factors^34^. Collectively these findings imply that tau regulates RNA and RBP function and the sequestration of RBPs in tau aggregates. Although some of the pathological interactions of tau have been identified from postmortem brain tissue, these studies in tissue are limited by their ability to capture transient interactions.

To overcome these limitations, recent in vitro interactome studies have leveraged proximity labeling as a tool to dissect both transient and stable interacting partners of proteins of interest using a bioengineered promiscuous biotin ligase^14, 35–37^. One such enzyme engineered from E.*coli*, TurboID, covalently binds biotin to lysine residues of proteins within a 10 nm range in minutes. IP methods limit the use of ionic detergents, such as those used in insoluble protein preparations, for identifying aggregate-prone proteins in disease.^38^ In contrast, the use of biotin leverages its strong affinity to streptavidin for enrichment of these biotinylated interacting proteins in strong, denaturing lysis conditions. Overall, proximity labeling has proved to be a useful tool in characterizing protein interactors across cell signaling pathways, aberrant interactions in disease models, and protein trafficking^37, 39–41^.

Here we applied a proximity labeling technique using split TurboID, a modified form of TurboID that activates only when two domains are in close proximity^41, 42^. We focused on the microtubule-binding repeat domain (MTBR) of tau, previously shown to form aggregate-like structures in human cell culture upon overexpression^43, 44^. Expression of two MTBR fragments carrying the N- and C-terminal of sTurbo yielded robust biotinylation, confirmed by immunoblotting and immunocytochemistry. Using affinity capture mass spectrometry, the sTurbo TauRD system captured both known tau-interacting proteins and predominantly nuclear proteins, many of which are involved in RNA binding and had not been previously identified as tau interactors. Many of the sTurbo-interacting RNA-binding proteins were also detected in post-mortem tissues and found to be enriched in the sarkosyl-insoluble fractions of AD brain tissues. Integrating these findings with independent tau interactome studies from AD cases and proteomic co-expression analysis in AD and related tauopathies further highlighted a strong correlation between NFT burden and RNA-binding proteins, many of which directly interact with tau in our sTurbo TauRD cell culture. Collectively, our study identified several novel tau-interacting proteins with relevance to human disease, providing promising candidates for mechanistic exploration.

## Results

### Establishment of a Split-TurboID TauRD Proximity Labeling System in Cell Culture

The microtubule-binding region (MTBR) forms the core of neurofibrillary tangles in the brain, which is a major pathological feature across tauopathies, including AD^45, 46^. Overexpression of the MTBR domain has been shown to induce aggregation in cell model systems^43, 44^. Here, we sought to develop a tau microtubule repeat domain (TauRD) split proximity labeling system to unbiasedly identify co-interacting partners associated with tau self-association and aggregation, similar to the well-characterized split-GFP TauRD HEK293 model systems^43, 47, 48^. As shown in **Fig. 1A**, we first generated two separate expression constructs, each encoding for 133 amino acids of the tau MTBR as the bait protein, fused to either the 8 kDa N-terminal split TurboID (N-sTurbo) or the 27 kDa C-terminal split TurboID (C-sTurbo) fragment. The N-sTurbo plasmid includes a V5 epitope tag, while the C-sTurbo is marked with an HA epitope tag. Upon co-expression, each construct is expected to produce a tau-MTBR that, when the two sTurbo biotin ligase fragments are brought into proximity, activates the biotin ligase enzyme, tagging proteins within a 10 nm range with intracellular biotin. This system enables the labeling of TauRD-interacting partners for downstream proteomic analysis, as depicted in the in vitro model (**Fig. 1B**).

**Figure 1.**
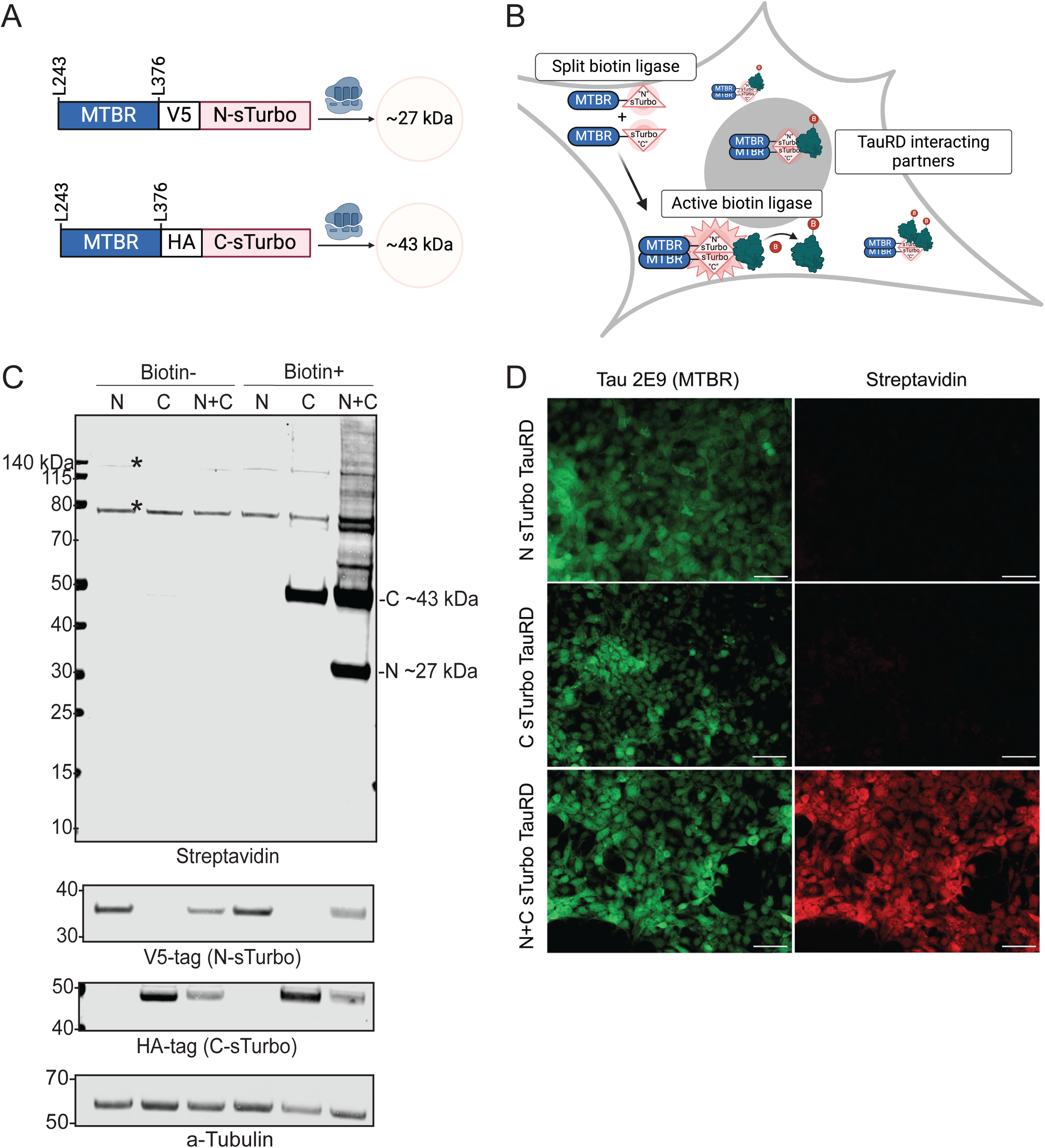
sTurbo TauRD expression and biotin ligase activity in a human cell line. **(A)** Design of split-Turbo Tau Repeat Domain (sTurbo TauRD) and respective anticipated molecular weights. The N-sTurbo is 8 kDa, while the C-sTurbo is 27 kDa; upon interacting, these recapitulate the functional activity of the complete 35 kDa E.*coli* biotin ligase. A unique biochemical tag was added to this design to distinguish the two split enzymes: V5 for N-sTurbo and HA for C-sTurbo. Each construct contains a 133 amino acid sequence of the four-repeat containing microtubule-binding domain (MTBR) from Tau 243-376, 441 numbering system, and houses a P301L pro- aggregation substitution. **(B)** Proposed model of sTurbo recombinant protein activity in vitro. Once the Tau MTBRs interact, the two sTurbo ligase fragments become functional and biotinylate proximal multimeric Tau protein interactors with or without the edition of 200 µM exogenous biotin. **(C)** Co-transfection with each split Turbo fragment displays robust biotinylation in the presence of exogenous biotin. sTurbo fragments were individually transfected (N or C) or co-transfected (N+C) into HEK293 cells. Groups were treated with serum-free media (Biotin-) or 200 µM biotin (Biotin+) for 1 hr. Streptavidin-conjugated dye was used to probe for biotinylated proteins. Biochemical tags (V5 or HA) confirmed the recombinant protein expression, and a-tubulin was probed for verification of equal protein loading across lysates. The C-sTurbo fragment displays low-level biotinylation, as it houses the ligase active site. WB revealed robust biotinylation in co-transfected lysates. Endogenously biotinylated protein bands are observed at ∼73, 75, and ∼130 kDa. **(D)** Immunocytochemistry (ICC) of respective N- and C-fragments confirms WB results of biotin ligase activity upon recombinant protein co-expression. Tau 2E9 MTBR-specific antibody detected recombinant protein expression while streptavidin dye captured biotinylated interacting partners. Scale bars represent 50 µm.

To assess expression and biotin ligase activity, we first expressed the N- and C-terminal sTurbo TauRD constructs alone or together in a HEK293 cell culture system. Protein lysates from untreated (Biotin-) and biotin-treated (Biotin+) cells were analyzed by Western blot (WB) for streptavidin and for the recombinant tags for the TauRD harboring the N- and C-terminal sTurbo proteins (**Fig. 1C**). N-sTurbo was detected at ∼27 kDa by blotting for the V5 tag in transfected lysates while the ∼43 kDa C-sTurbo TauRD fragment was detected by blotting for HA. In the absence of biotin, only endogenously biotinylated proteins were observed at ∼135 and ∼75 kDa. However, upon biotin treatment, we detected self-biotinylation of the C-sTurbo TauRD fragment at ∼43 kDa in cells expressing just the C-sTurbo TauRD recombinant protein. This partial biotin activity is consistent with prior studies^41, 42^. In contrast, robust biotinylation of both the C-sTurbo and N-sTurbo TauRD fragments (∼43 and ∼27 kDa) was observed in cells only after co-expressing both constructs and treated with biotin. Furthermore, extensive protein biotinylation was detected across a wide molecular weight range with a streptavidin infrared label, indicating effective labeling of not only the sTurbo TauRD fragments, but also potential interacting proteins. In a complementary approach, immunocytochemistry (ICC) was also performed for cells expressing either the N- or C-sTurbo TauRD fragments alone or in combination. The sTurbo TauRD fragments were detected with a specific MTBR-tau antibody (2E9), while biotinylated proteins were visualized with a streptavidin fluorescent dye. Imaging did not display biotinylation when the TauRD fragments were expressed alone; however, robust biotinylation was clearly seen upon co-transfection (**Fig. 1D**). The absence of detectable biotinylation by ICC for C-sTurbo tau alone, compared to detection by WB, may be due to differences in assay sensitivity. Collectively, results from both WB and ICC analyses confirm that the sTurbo TauRD proximity labeling system successfully labels tau interacting partners when both N- and C-sTurbo fragments are co-expressed, with minimal self-biotinylation occurring when fragments are transfected separately.

### Development and Validation of a Bicistronic sTurbo TauRD Construct for Uniform Expression and Functional Proximity Labeling in HEK293 Cells

To overcome non-uniform co-transfection and, consequently, differences in gene expression within and across experimental groups, we generated a bicistronic plasmid containing both N- and C-sTurbo TauRD fragments separated by a 2A self-cleaving linker^49^. As illustrated, the 2A linker was inserted after the N-sTurbo and before the subsequent MTBR domain (**Fig. 2A**). Biotin ligase expression and activity in HEK293 cells transfected with single sTurbo TauRD bicistronic plasmid was confirmed by WB. The N- and C-sTurbo biotinylated TauRD fragments were observed at their expected molecular weights, ∼27 and ∼43 kDa, respectively, whereas transfected cells only displayed endogenously biotinylated protein bands at ∼135, 78, and 75 kDa (**Fig. 2B**). Additionally, a band at ∼75 kDa reacted with the V5 antibody, but was not detected by HA (**Fig. S1A**). Prior reports have described a potential overabundance bias toward the first coding sequence of some 2A linkers^50^. To address this, a semi-quantitative WB was used to measure the biotinylated N- and C-sTurbo TauRD protein signal intensities across four replicates. The N- to C-terminal ratios were approximately 1:1, indicating minimal labeling bias between fragments due to uncleaved 2A (**Fig. S1B**). To assess localization and biotinylation activity of the recombinant proteins, we performed ICC. sTurbo TauRD localized throughout both the nucleus and cytoplasm, often forming speckle-like inclusions in the cytoplasm^51^. These inclusions largely overlapped with the biotinylation signal, likely due to tau self-biotinylation and biotin-labeled proximal interactors (**Fig. 2C**). Overall, these results confirm the successful expression and activity of sTurbo TauRD in HEK293 cells using a single bicistronic plasmid containing both N- and C-sTurbo TauRD. These results also reveal the labeling of potential co-aggregating partners across a range of molecular weights.

**Figure 2.**
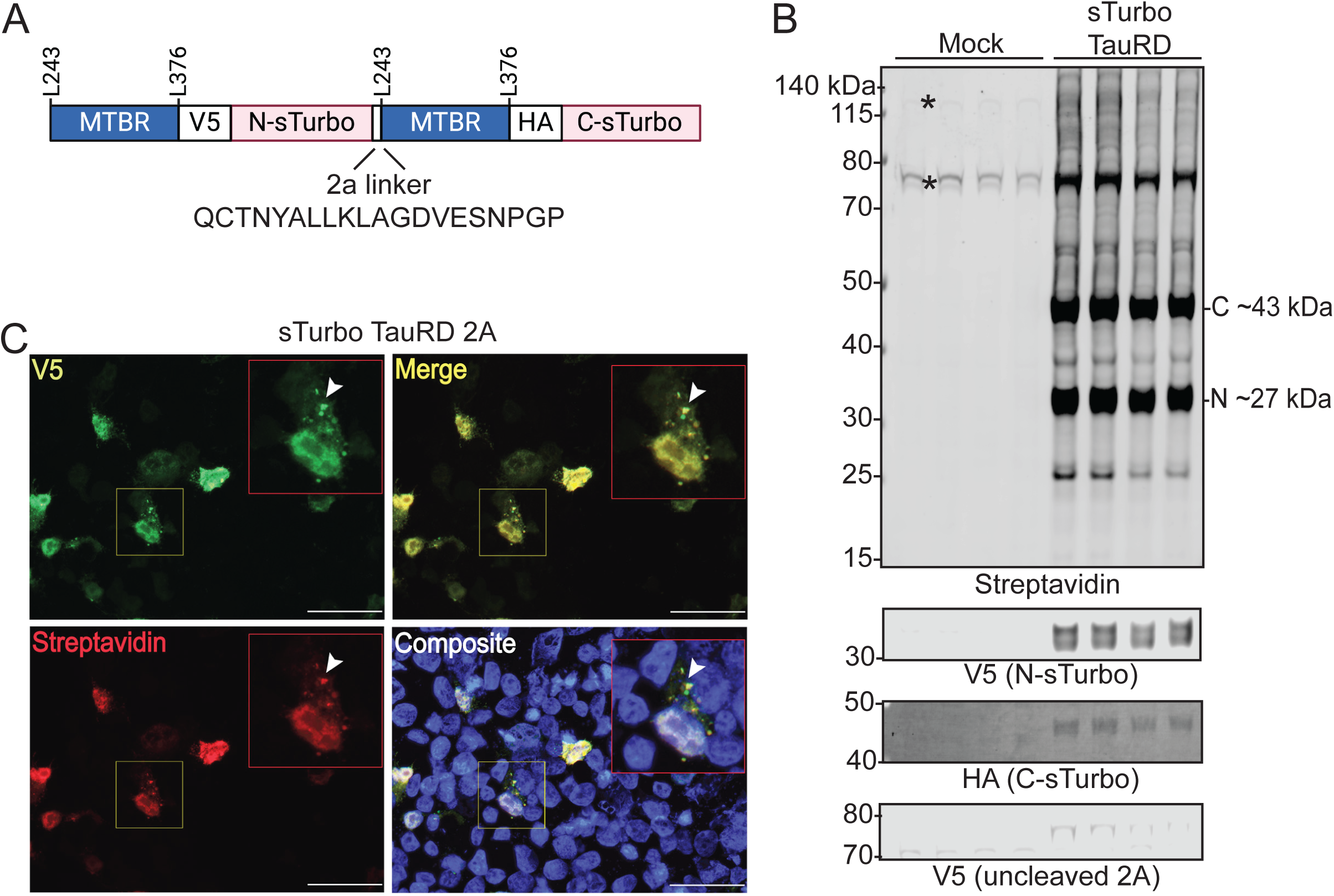
Development and characterization of a bicistronic sTurbo TauRD construct and functional proximity labeling. **(A)** Design of split ligase construct utilizing a single 2A plasmid to mitigate co-transfection efficiency variations. An e2A self-cleaving linker enables a 1:1 stoichiometric expression of each ligase fragment in the same cell. **(B)** Confirmation of biotin ligase expression and activity in HEK293 cells transfected with a single plasmid sTurbo TauRD via WB analysis. The N- and C-sTurbo fragments exhibit reactivity to their respective biochemical tags and demonstrate self-biotinylation with only a weak proportion of uncleaved recombinant protein demonstrated at ∼75 kDa in the V5 channel. Transfection reagents without plasmid (Mock) reveal endogenously biotinylated proteins at ∼130, ∼75, and ∼73 kDa. **(C)** ICC displays Tau-positive inclusions in the cytoplasm and nucleus, co-localizing with biotinylated proteins (yellow). Rabbit anti-V5 tag (green) targets the sTurbo TauRD recombinant protein, while streptavidin dye (red) portrays biotinylated proteins. Images were captured at 40X magnification. Scale bars represent 50 µm.

### Proximity Labeling of Tau MTBR Interactors and Mass Spectrometry Analysis in Cells Reveals Established and Novel Tau-associated Pathways

Antibody-based affinity purification coupled to mass spectrometry (AP-MS) studies have provided a breadth of typical and pathophysiological interactors of tau from both cells and tissues, including RNA-binding proteins (RBPs), 14-3-3 binding proteins, numerous cytoskeleton components, heat-shock proteins, and proteasome subunits^13–16^. Here, we sought to unbiasedly identify interacting proteins of the tau MTBR using our sTurbo expression system. To this end, HEK293 cells (n=10 biological replicates) underwent a standard streptavidin affinity purification (SA-AP) and label-free quantitative mass spectrometry (LFQ-MS) workflow, as previously described^36, 39^ and outlined in **Fig. 3A**. As expected, sTurbo TauRD lysates showed consistent and robust biotinylation across replicates, including self-biotinylation of sTurbo TauRD fragments **(Fig. S2A)**. To confirm pulldown upon SA-AP, gel electrophoresis and silver staining was performed from each eluent, which confirmed the enrichment of proteins in sTurbo TauRD samples as well as non-specific protein binding to streptavidin beads in the mock samples, that were omitted from further investigation **(Fig. S2 B-C)**. Following WB and silver stain quality control assays, Mock (n=6) and sTurbo TauRD (n=10) pulldowns were digested directly on beads for downstream proteomic analysis^36, 39^. Following LFQ-MS, resulting intensities advanced through an in-house AP-MS pipeline, including quality control and imputation for missing values^35, 36, 39^ as described in the methods **(Supplemental Tables 1-3)**. Unique MTBR peptides, along with those corresponding to the custom N and C-sTurbo FASTA sequences were also identified **(Supplemental Table 4)** which confirmed biotinylation and pulldown of the expected TauRD domains. Overall, 2,167 proteins were retained for subsequent differential enrichment analyses (DEA).

**Figure 3.**
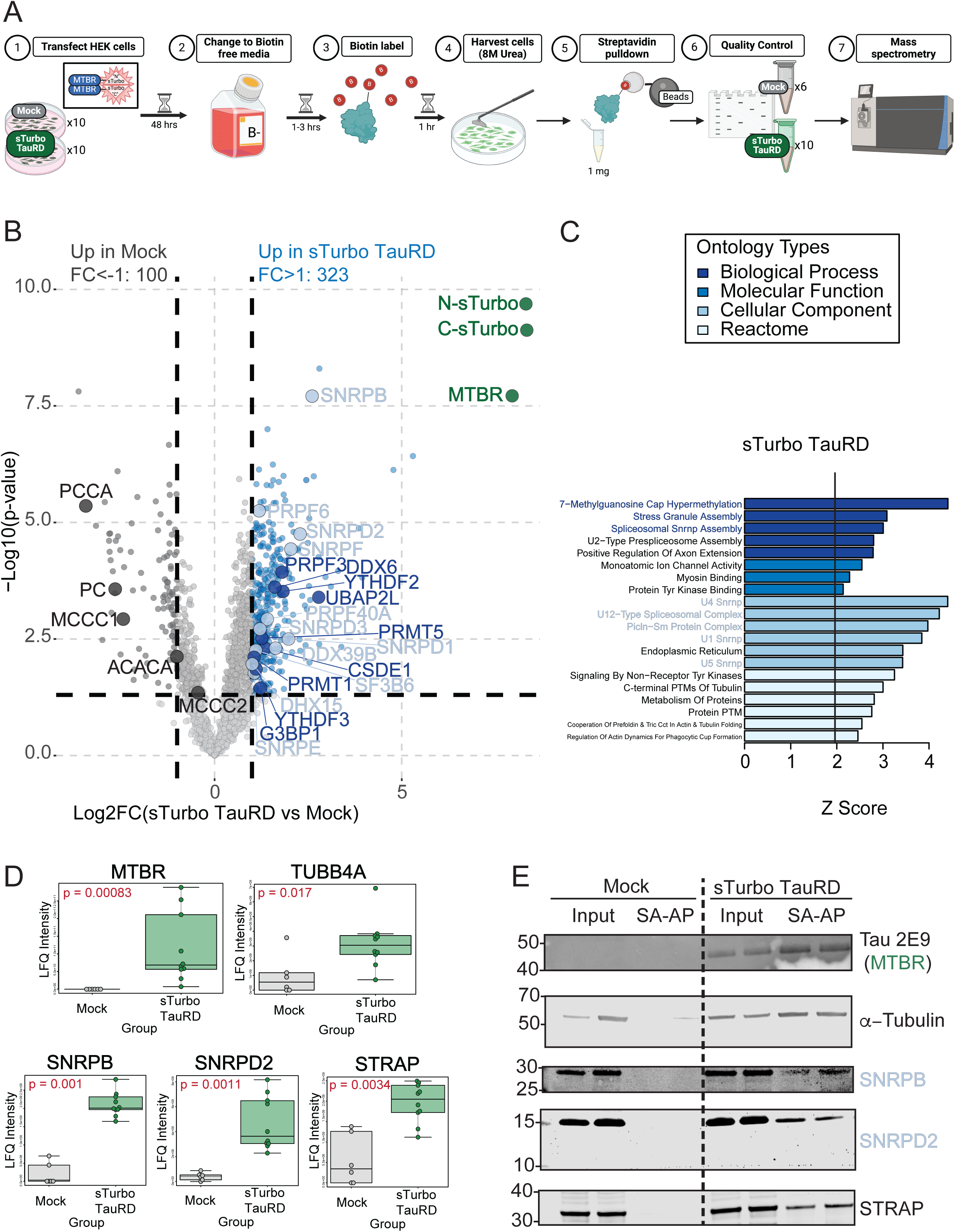
Differential abundance analysis of TauRD proteomics demonstrates nuclear protein enrichment. **(A)** Schematic experimental workflow for identifying protein-protein interactors of the MTBR domain of Tau. Ten replicates of HEK293 cells across three batches were transfected, underwent a biotin washout period, labeled with 200 µM biotin for one hour, and harvested in 8M Urea with protease inhibitors. 1 mg of total protein from whole cell lysate was incubated with streptavidin-conjugated magnetic beads for 1 hour to enrich for biotinylated protein interactors. Samples underwent quality control by total protein silver stain. Biotinylated proteins were investigated via mass spectrometry. **(B)** Mass spectrometry identified over 700 significant proteins in sTurbo TauRD lysates compared to controls. Volcano plot displays the Log2 Fold Change between groups plotted on the x-axis while –Log10 unadjusted p-value, as determined by one-way ANOVA, is plotted on the y-axis. A p < 0.05 (-Log10 = 1.3) and a 2-fold change higher than control (Log2FC = +/-1) cutoff was employed for stringency to identify potential candidate proteins for validation. Blue points indicate proteins significantly enriched in the sTurbo TauRD samples (n=323), while dark grey dots indicate proteins enriched in the Mock samples (n=100). Green points represent overexpressed exogenous proteins with the highest observed fold change and significance. Mock-enriched proteins are related to endogenously biotinylated carboxylases, labeled in the plot, and contaminating keratin proteins. **(C)** Top Gene Ontology (GO) terms for these putative TauRD interacting proteins (n=323) confirmed various pathways, including cytoskeletal components, RNA binding, and protein post-translational modifications (PTMs)**. (D)** Canonical TauRD interacting proteins, such as the Tau-MTBR itself and a-Tubulin, were selected for validation and served as positive controls. Candidate RNA-binding proteins SNRPB, SNRPD2, and STRAP are significantly enriched in the sTurbo group as visualized via boxplots of raw LFQ intensities between Mock and sTurbo TauRD samples. NA values were converted to zero for visualization. Boxes represented median of each group with 25^th^ and 75^th^ percentiles. P-values were generated by Kruskal-Wallis test. **(E)** Candidate proteins were additionally validated through WB. Total lysate input is equal across both groups. Only the sTurbo Tau-expressing samples maintained interactions with these proteins upon streptavidin affinity purification.

DEA revealed 1090 total significantly different proteins; 761 were increased in sTurbo TauRD pulldowns whereas 329 proteins were enriched in Mock pulldowns. As anticipated, the over-expressed recombinant sTurboID N- and C-terminal specific proteins and the core TauRD domain (shared between the N and C- term fragments) were most significantly enriched in pulldowns with Log_2_ fold change (FC) of 8.3, in both fragments with a false discovery rate (FDR) corrected p-value of 8.12E-07 in C-sTurbo and 4.44E-07 in N-sTurbo fragments. Importantly, the similarity of FC enrichment across the fragments verifies the 1:1 expression ratio of the 2A products. Consistently, the TauRD which is translated contiguously with the N- or C-term sTurboID fragments had a log_2_ fold change FC of 7.95 (FDR: 7.04E-06) **(Supplemental Tables 5 & 6)**. We applied a two-fold change cut-off threshold (Log_2_FC ≥1) to identify enriched proteins and conduct Gene Ontology (GO) analysis. With this thresholding in place, sTurbo TauRD resulted in 323 enriched proteins, while 100 were increased in Mock samples **(Fig. 3B)**. GO analysis of Mock enriched proteins was associated with vitamin binding (biotin metabolism) and skin epidermis (keratins) consistent with enrichment of endogenously biotinylated proteins as would be expected in Mock samples. In contrast, the proteins enriched by sTurbo TauRD included the MTBR fusion proteins, as well as other proteins with known tau biology, including interactions with microtubule subunits and an over-representation of RNA-binding proteins associated with stress granules (G3BP1, CSDE1, DDX6, UBAP2L, and YTHDF3) and the spliceosome complex^15, 16, 21, 22, 25, 27, 29, 30, 52–57^. Additionally, sTurbo TauRD GO terms reflected 7-methylguanosine cap hypermethylation and Tyrosine (Tyr) kinase binding **(Fig. 3C, Supplemental Table 7)**.

Protein levels were visualized by volcano plot and were significantly enriched in the sTurbo TauRD interactome over Mock samples (**Fig. 3D**). To confirm the enrichment of biotinylated proteins and validate the mass spectrometry results, WB was performed on several candidate proteins following sTurbo tau MTBR pull- downs. This included canonical tau-interacting proteins, such as Tau-MTBR (detected using the Tau2E9 antibody) and α-tubulin^3, 4, 58, 59^, to compare enrichment levels across input lysates and pulldown eluents. In the sTurbo TauRD SA-AP pulldown, RNA binding proteins such as SNRPB, SNRPD2, and serine/threonine kinase receptor-associated protein (STRAP) were enriched in the sTurbo TauRD lysates, but not controls, consistent with the individual MS findings (**Fig. 3E)**.

To assess the disease relevance of our in vitro tau interactors to human brain tau interactors, our sTurbo TauRD interactors were compared to tau interactome datasets obtained from antibody-based AP-MS studies using Tau antibodies (Tau-5, total tau; PHF-1, phosphorylated pathological Tau, **Supplemental Table 8**) from AD frontal cortex^15, 16^. Overall, 128 proteins were consistently enriched across sTurbo TauRD and the complementary AP-MS human Tau interactomes (**Fig. 4)**. Unsurprisingly, tau protein binding was among the top GO terms, alongside secretory granule binding, unfolded protein binding^60, 61^ and clathrin adaptor activity^62, 63^. Meanwhile, brain-derived AP-MS tau interactomes shared proteins relating to synaptic plasticity regulation^64–68^, calmodulin binding^69, 70^, and ATP-dependent activity^71, 72^ **(Fig. 4A)**. sTurbo TauRD identified many distinct interacting partners and was uniquely enriched in RBPs relating to nuclear localization sequence binding and nuclear import **(Fig. 4).** In summary, this proteomic assay reveals both known and novel interactors of tau-MTBR, reflecting shared tau biological functions across previously identified human tau interacting partners and model systems.

**Figure 4.**
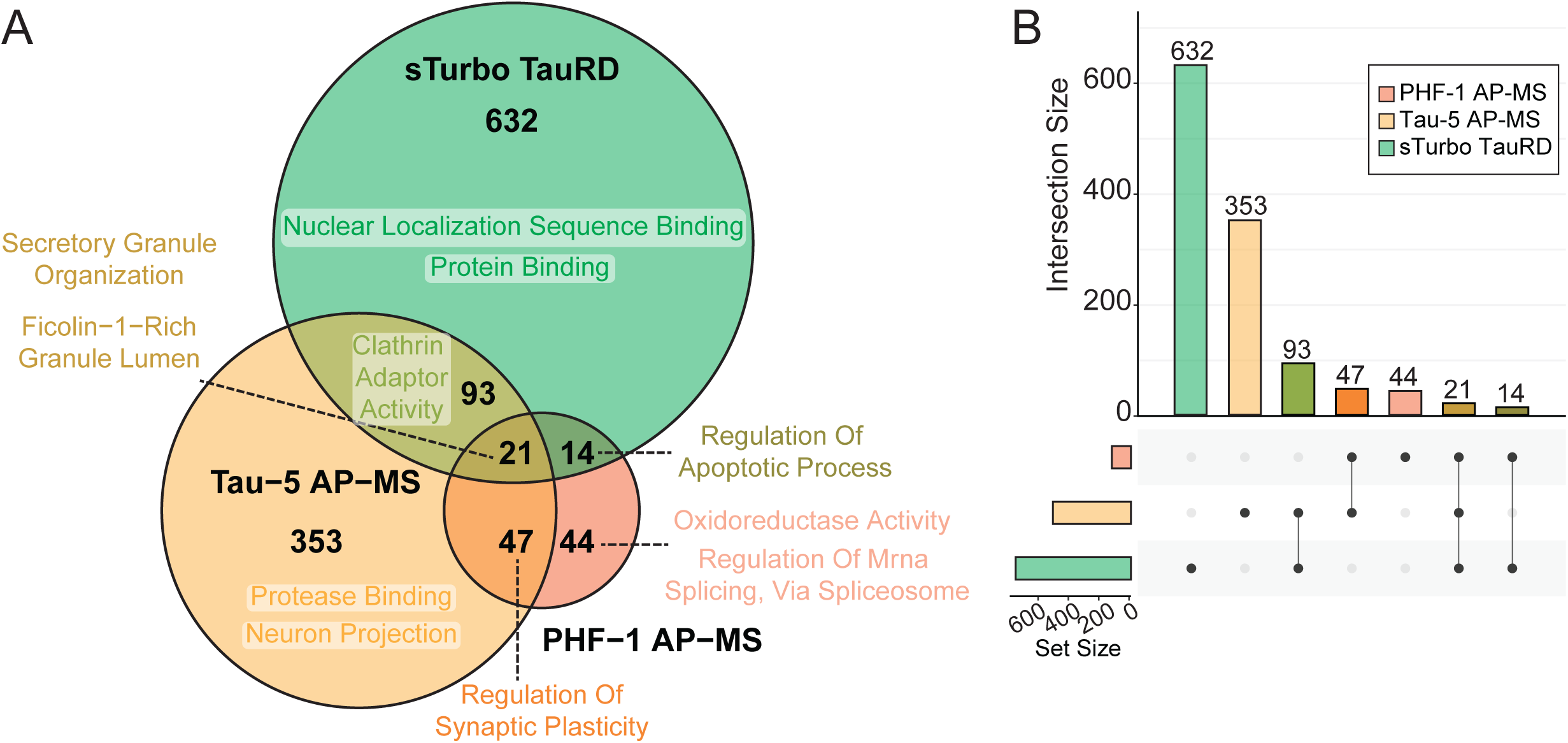
Intersection of tau-interactome datasets reveals core protein functions shared between human AD Tau and sTurbo TauRD. **(A)** Venn Diagram of enriched proteins compared to negative controls across sTurbo TauRD, combined Tau AP-MS, and frontal cortex proteomics across Tauopathies and top GO terms. **(B)** UpSet plot of resulting intersecting dataset proteins.

### Network Analysis of Human AD and Progressive Supranuclear Palsy Brain Tissue Identifies Protein Modules Associated with Tau Pathology

Interacting proteins identified through biotinylation enrichment may have been insoluble or transiently associated with pathological proteins,^73^ and therefore missed in human interactome datasets generated by standard IP. Consequently, proteins exclusively enriched in the sTurbo TauRD proteomics may still show disease relevance. To further understand the association of the TauRD interacting partners to disease, we integrated the tau interactome data with previously generated proteomics data from human post-mortem bulk frontal cortex samples^74^, including control (CTL, n=46), Alzheimer’s disease (AD, n=49), and progressive supranuclear palsy (PSP, n=26) **(Fig. 5A, Supplemental Table 9)**. After data quality control, this tandem mass tag (TMT) mass spectrometry-based human dataset consisting of 9,641 proteins was analyzed using established pipelines for systems-level network analysis^18, 75, 76^ **(Supplemental Table 10)**.

**Figure 5.**
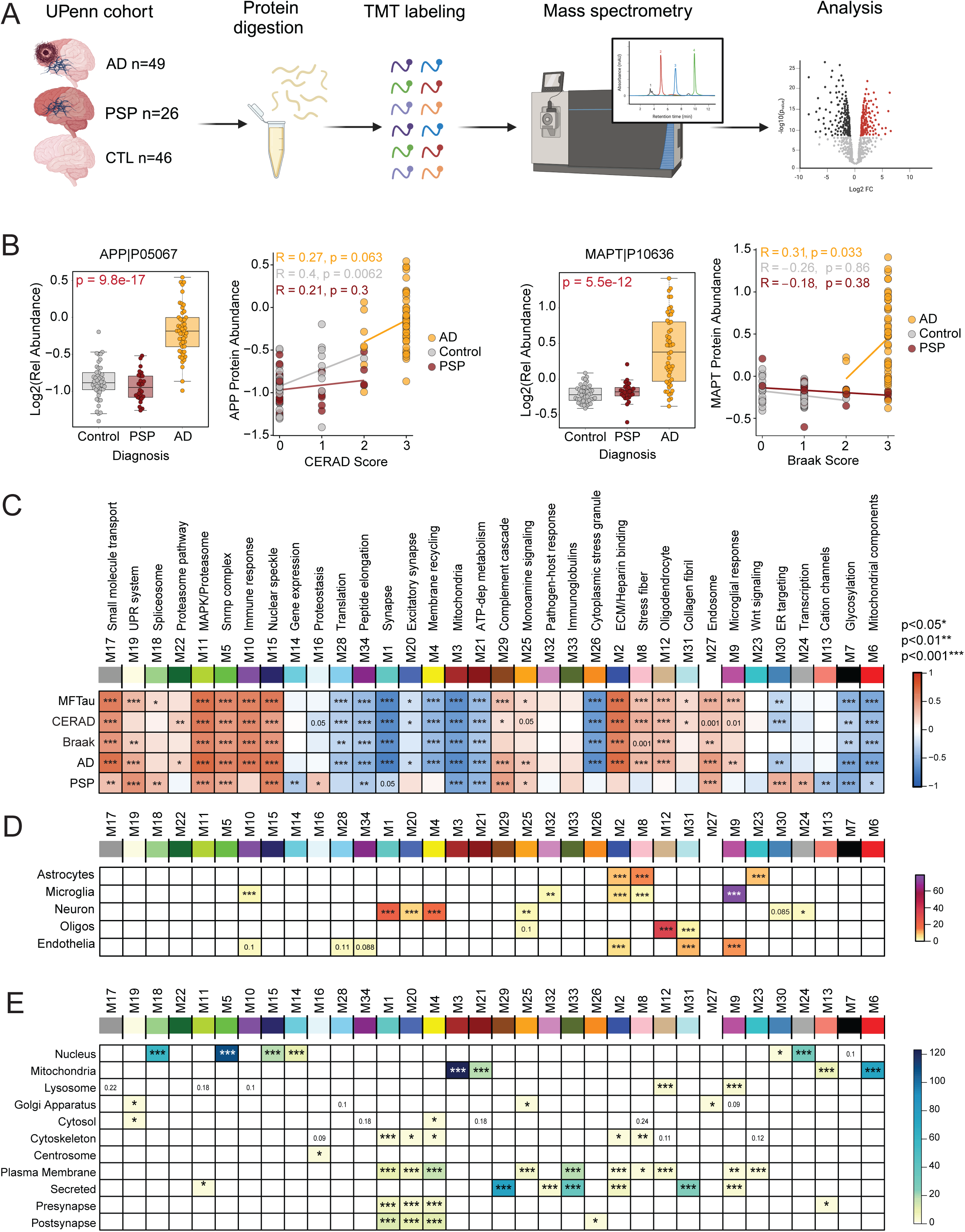
Network analysis of human AD and progressive supranuclear palsy (PSP) brain tissue. **(A)** Schematic of University of Pennsylvania (UPenn) brain tissue selection and experimental overview. AD (n=49), PSP (n=26), and CTL (n=46) frontal lobe tissues underwent proteolytic digestion, TMT labeling, and mass spectrometry. A relative protein abundance matrix was used for downstream analysis. **(B)** Quality control measures (CERAD and Braak) across PSP and AD groupings. APP levels across groups are related to increased amyloid burden (CERAD) in AD cases but not PSP. Similarly, MAPT levels are highest in AD and correlate to higher Braak staging in disease. PSP cases do not display increased amyloid deposition or frontal lobe Tau burden. **(C)** AD and PSP proteins (n=9,661) were analyzed via Weighted Gene Network Analysis (WGCNA) and separated into 34 highly related protein co-abundance modules. Each module name is summarized from significant Gene Ontology (GO) terms. Those highly related to Alzheimer’s Disease include M2 (ECM/heparin-binding) and M5/M15 (Snrnp complex, nuclear speckle). Signed bicor r-value of module abundance to Tau pathology (MFTau, Braak), Amyloid burden (CERAD) and diagnosis. Red represents a positive correlation, while blue describes a negative correlation (*, p<0.05; **, p<0.01; ***, p<0.001). **(D)** Neural cell type marker enrichment by Fisher’s Exact Test (FET) to modules. FDR corrected, Benjamini-Hochberg (BH) –Log10(p-values visualized. **(E)** Subcellular localization marker enrichment by FET to modules. -Log10(FDR, BH p-values) visualized.

As expected, Amyloid precursor protein (APP), which in bulk AD brain proteomic datasets has been previously shown to be a proxy for β-amyloid (Aβ) deposition^77^, was significantly higher in AD cases than controls and PSP. Elevated APP levels also positively correlated to neuropathological Aβ burden (CERAD score). However, as expected, APP levels between control and PSP cases, did not show any differences in relative abundance^76, 77^. Braak staging, a metric characterizing both the severity and localization of pathological tau spread throughout the brain, was significantly elevated in AD. MAPT protein abundances also correlated to pathological tau burden (Braak stage)^78^. Meanwhile, this change was not observed in PSP or control cases **(Fig. 5B)**. While PSP also exhibits tau tangle pathology, the highest burden remains in the temporal lobe rather than the frontal lobe region analyzed by TMT-MS based proteomics.

To gain more insight into system-level changes correlating to disease and pathology, a weighted gene co-expression analysis (WGCNA) pipeline for network analysis was applied to ascertain modules of highly co-expressed proteins across all tissues^79^. This approach resulted in 34 modules, with M1 being the largest (n=881 proteins) and the smallest being M34 (n=35) **(Supplemental Table 11)**. Significant biological processes for each module were identified through GO analysis and named to summarize a representative ontology or cell type involvement (**Supplemental Table 12).** Correlation analysis of key pathological traits, including immunological tau values (Mid-frontal, MFTau), CERAD score, Braak staging, and diagnosis of AD or PSP, reveal significant modules relating to disease processes **(Figure 5C).**

Modules uniquely increased in AD were enriched in inflammatory brain immune protein markers, including those specifically expressed in astrocytes, microglia, and endothelia (M10, M2, M8 and M9) **(Fig. 5D, Supplemental Table 13)** consistent with previous reports^18^. Many of the AD associated modules also corresponded to proteins localized to the ‘Golgi Apparatus’, ‘Cytoplasm’ and ‘Plasma Membrane’ (**Fig. 5E, Supplemental Table 14)**. Modules positively correlating to PSP were enriched in nuclear proteins (M18, M5, M15, M30, M24). Consistent with prior reports^17, 21, 32, 57, 80–83^ of shared biology across different Tauopathies, M5, M15, and M18 enriched in nuclear proteins were increased in both AD and PSP. Overall, we identified several modules that positively correlate with both AD and PSP and are associated with neuropathological tau burden. However, due to the limitations of bulk cortex proteomics, it remains unclear whether the proteins within these modules directly interact with tau.

### Integrating the TauRD Interactome with AD and PSP Network Modules to Identify Disease-Associated Modules Linked to Tau PPIs

To assess whether tau interactors are enriched in disease-associated protein modules in human AD and PSP, we integrated the recombinant sTURBO TauRD and human tau interactomes with the cortical AD and PSP networks. To this end, a hypergeometric Fisher’s exact test (FET) was used to identify modules in the human AD and PSP co-expression network that were significantly enriched with tau interactors from the sTURBO TauRD and human tau interactome datasets. **(Fig. 6A, Supplemental Table 15)**. Six modules exhibited significant overlap with the tau interactomes. Namely, human AD brain tau interactors showed significant overlap with two sTurbo TauRD enriched modules (M16, Proteostasis; M28, Translation) as well as enrichment in M22 (Proteasome pathway) and M11 (Proteasome/MAPK signaling) **(Fig. 6B)**. As expected, several tau interactors from human brain tissue were uniquely enriched in neuronal and synaptic modules containing proteins that are not strongly expressed in HEK293 cells **(Fig. 6 B-C)**. This analysis also revealed seven modules (M17, M19, M18, M5, M10, M16, and M28) encompassing 256 proteins significantly enriched in TauRD interactors. Of these, five modules (M17, M19, M18, M5, M10) were enriched in TauRD interactors, but did not display overrepresentation of human tau interactors. Because conventional antibody-based IP cannot readily affinity-capture insoluble protein interactions^38^ we hypothesized this may be due to the insoluble nature of module proteins. We further examined whether the sTurbo TauRD interactors we identified were enriched in the insoluble human AD proteome using previously published data **(Supplemental Tables 16 & 17)**^84^. This analysis identified several disease-associated modules, specifically M5, M15, and M18, that overlapped with the insoluble AD proteome **(Fig. 6C)**. These modules include proteins such as hnRNPs (HNRNPA2B1, TARDBP)^83, 85^ and spliceosome members (SNRPD1, SNRPD2, SNRPE)^56^. Notably, M5, M15, and M18 were significantly enriched in RBPs localizing across the nucleus, nuclear speckles, and cytoplasm. These modules relate to splicing regulation function and RBPs containing canonical RNA Recognition Motif (RRM) domains (**Fig. S3 A-C)**. M5 and M18 have the most similarity across RBP types: viral RNA regulation, spliceosome, RNA stability & decay, and nuclear export and translation regulation functions are all represented. These modules also slightly differ in types of enriched RBPs. M5 houses RBPs uniquely mapping to nucleolus, M15 is the only disease-related insoluble nuclear module with Zinc Finger (ZnF) domain-containing proteins represented, and M18 RBPs are also involved in 3’ end processing of pre-mRNA^86^. Overall, the sTurbo TauRD interactome overlapped with three tauopathy-associated modules that were not enriched with human tau interactors, likely in part due to the enrichment of insoluble nuclear proteins relating to histone and DNA-binding, nuclear transport, polyadenylation factors, and splicing.

**Figure 6.**
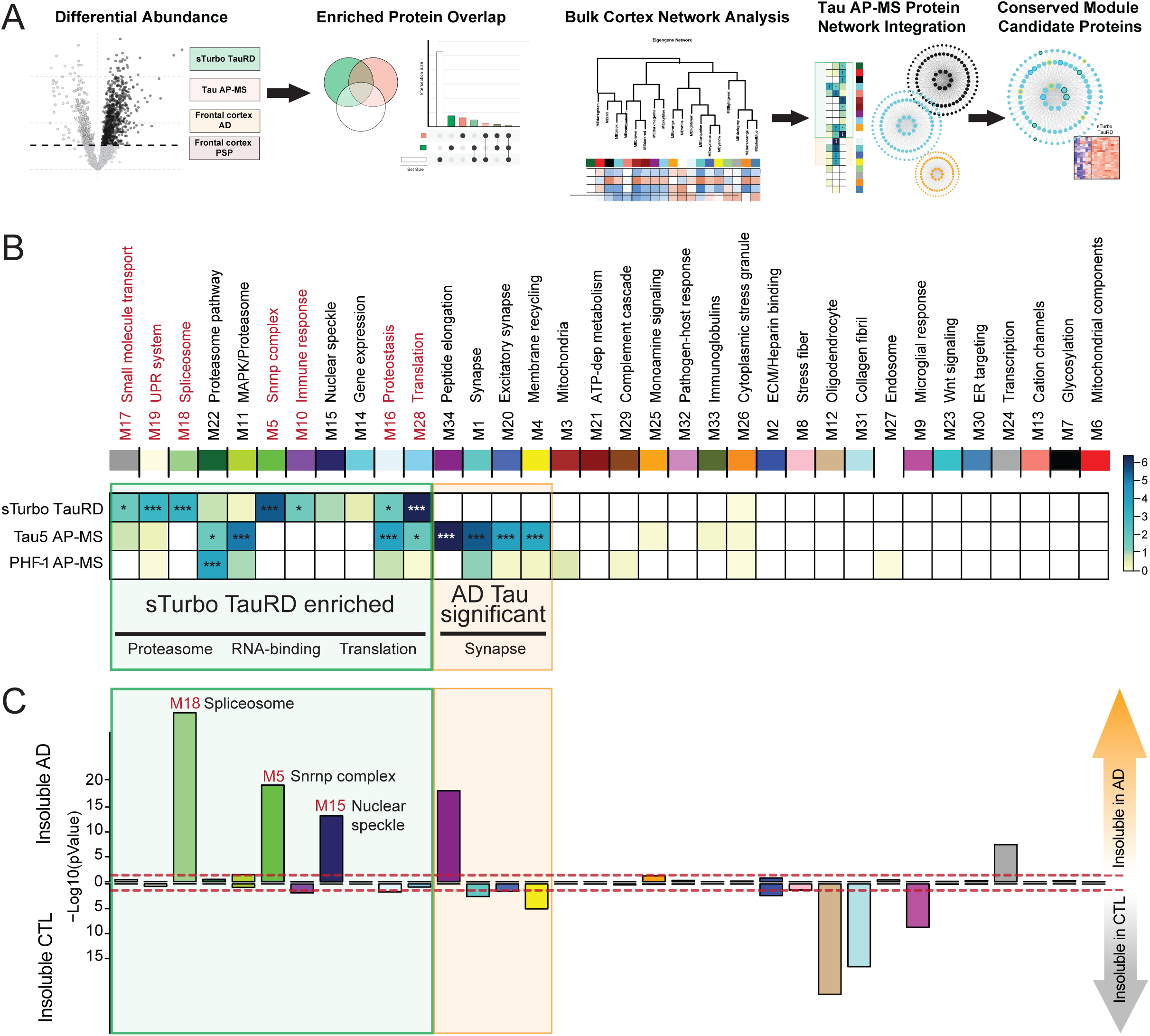
Integration of the TauRD interactome with a human tauopathy proteomic network identifies RNA-binding protein modules associated with AD and PSP that are enriched in insoluble proteins. **(A)** Overview of data integration and visualization steps. A differential abundance of proteins in TauRD (green), Tau AP-MS (Tau5 and PHF-1 antibodies; orange) and human AD & PSP bulk frontal cortex proteomics from Tauopathies (white) datasets was conducted. Total protein overlap was visualized by Venn Diagram and UpSet plots for comparison of enriched GO terms of intersecting groups. Then, enriched Tau AP-MS studies (green & orange) were integrated into the human AD & PSP bulk frontal cortex proteomics network to determine protein overlap across modules. Tau AP-MS overlapping proteins were also visualized by their degree of module membership in modules highly correlating to disease. **(B)** FET of enriched protein overlaps with tauopathy brain network modules across sTurbo TauRD and AD Tau interactomes, shown within a green square. 7 modules were significantly enriched in sTurbo TauRD enriched proteins. Disease-associated proteosome modules M22 and M11 were also reflected in AD Tau AP-MS interactomes with trends toward significance in sTurbo TauRD. Most significant AD Tau interactome modules discordant to sTurbo TauRD (M1, M20, & M4; highlighted with an orange square) were enriched in synaptic proteins. FDR corrected, Benjamini-Hochberg (BH) –Log10(p-values) visualized. **(C)** FET of brain network modules to the insoluble proteome from TMT-labeled AD brain and CTL brain lysates reveals modules related to sTurbo TauRD are generally insoluble protein enriched and related to proteosome, RNA-binding, and translation ontologies. –Log10(uncorrected p-values) visualized.

To better understand the intersection between the detergent-insoluble proteome in tauopathies and tau interacting proteins mapping across the three insoluble RBP-rich disease-associated modules, the signal intensities of the sTurbo TauRD interactome and top 150 correlated proteins in each module were visualized **(Fig. 7A and Fig. S4)**. Several proteins intersected across all three tau interactome datasets in M5 (SNRNPD3, HNRNPA2B1, and RBM25) and M18 (SNRPD2, HNRNPK, and EDC4) while sTurbo TauRD interactors were represented across both soluble and insoluble hub proteins in these selected modules **(Fig. 7 B-D).** Surprisingly, human AD tau-interacting proteins predominantly overlapped with insoluble proteins represented in modules M15 and M18 (**Fig. 7 and Fig. S4**). In contrast, sTurbo TauRD primarily captured proteins correlating with Braak staging across three modules. Notably, these included CHD4 and MACROH2A1 in M5, WDR82 and PAPOLA in M15, and SF3B6 and SAFB in M18 (**Fig. 7 E-G**), highlighting their potential relevance to disease. Although these proteins have been associated with models of AD^87–92^, they have not previously been identified as tau interactors. Here, we demonstrate the utility of integrating sTurbo TauRD interacting proteins with a tauopathy proteomics network encompassing both AD and PSP cases to identify disease-relevant modules enriched with tau interactors. This analysis revealed several RNA-binding protein modules that were enriched with TauRD-interacting proteins and positively associated with tau pathology that were likely missed in human tau interactome datasets due to their transient or insoluble nature in the human brain.

**Figure 7.**
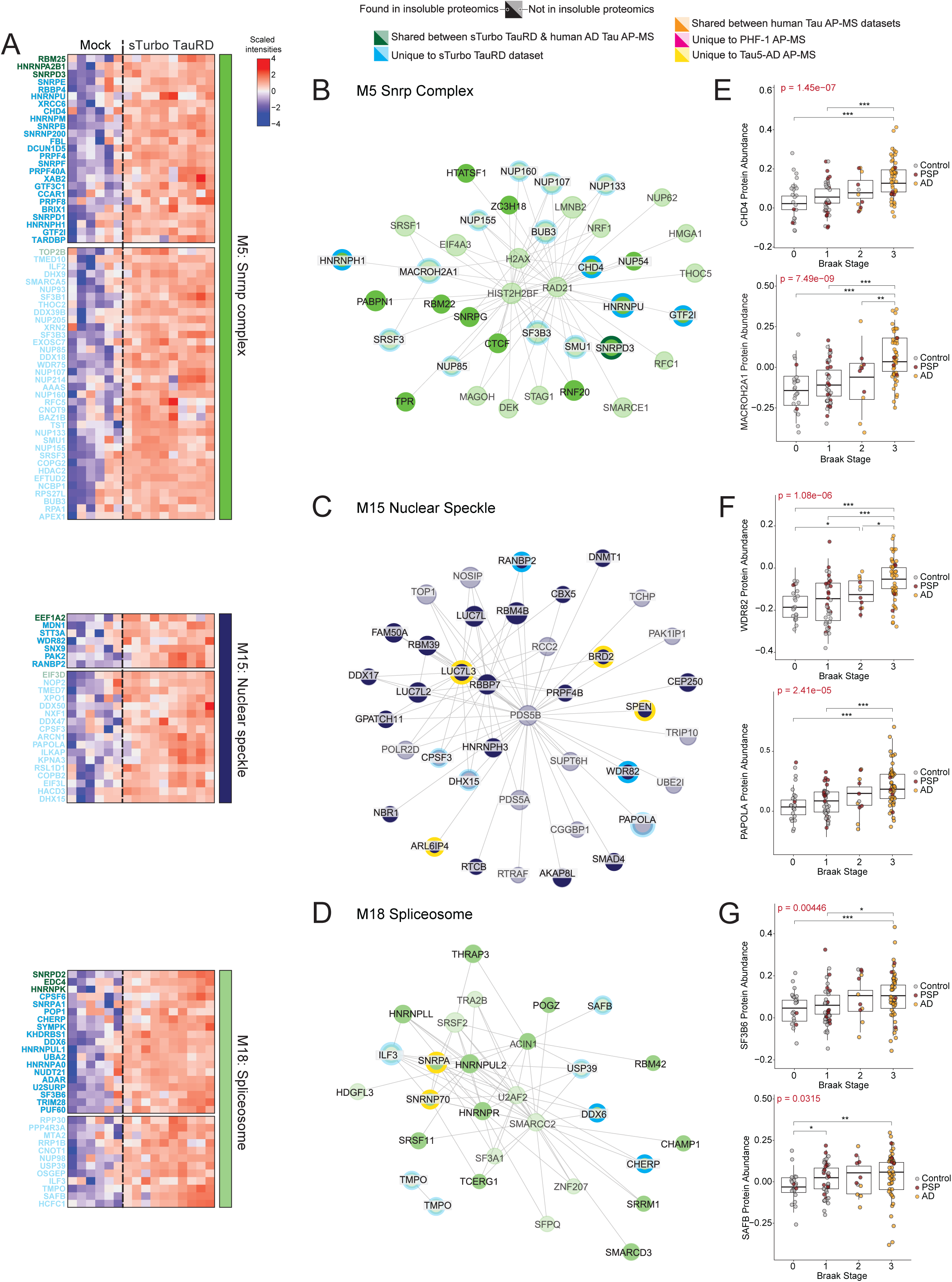
Network visualization of significant human RNA binding protein modules that overlap with the sTurbo TauRD interactome. **(A)** Heatmap of sTurbo TauRD significant FET modules enriched in RBPs. LFQ data was intersected with an RBP comprehensive database (Embl) to select module RBPs. Intensity values across sTurbo TauRD and Mock individual samples were z-scored and scaled for visualization. Greater intensities are colored in hues of red, whereas purple represents lower protein intensities. Protein IDs are colored according to intersection across Tau interactome datasets: Tau interactome proteins are identified within the module, either from sTurbo TauRD (unique, blue; shared, green) or described in human AD Tau AP-MS (warm colors). Proteins mapping to insoluble proteomics data are shown opaquer. **(B)** Fructerman-Reingold projection of module hub proteins connected by the top 5% of Topological Overlap Matrix (TOM) values. Edge weights are representative of the degree of relatedness of the node to the hub proteins. Protein nodes are colored as described above. Visualization for M5 Snrnp complex, **(C)** M15 Nuclear speckle, **and (D)** M18 Spliceosome for disease-related RBP-rich modules enriched **(E-G)** sTurbo TauRD unique proteins (M5:CHD4, MACROH2A1; M15:WDR82, PAPOLA; M18: SF3B6, SAFB) correspond to Braak staging. Protein intensities were plotted across human AD & PSP bulk frontal cortex Braak stages. A Kruskal-Wallis rank sum is displayed across all Braak stages while a Student’s t-test was implemented across individual comparison levels and significance denoted (*, p<0.05; **, p<0.01; ***, p<0.001).

## Discussion

In this study, we demonstrate that the in vitro TauRD interactome identifies conserved cellular machinery associated with tubulin binding and spliceosomal proteins. By integrating with additional tau interactome studies and bulk brain proteomics, we show that the TauRD interactome provides shared insight into protein-protein interaction (PPI) changes, further reinforcing the role of RNA-binding proteins (RBPs) in AD and related Tauopathies^17, 20, 21, 25, 30, 32–34, 56, 57, 80, 93^. We uncovered new tau interacting proteins by comparing TauRD proteins from bulk frontal cortex insoluble proteomes. These protein abundances also increase with Braak staging, complementing previous work demonstrating RBPs aggregate alongside neuropathological tau burden^56^. The TauRD interacting proteins within disease-related protein modules were enriched in nuclear pathways consistent with prior studies highlighting the relevance of splicing factors and RBPs in AD pathogenesis^16, 94^ and loss of function with downstream splicing defects^16, 17, 22, 32, 56, 80^. Notably, the aggregation of these RBPs in AD also follows tau burden by Braak staging^56^, suggesting a potential connection between pathological tau spread and the RBP and splicing dysregulation observed in tauopathies.

Previous work by our group and others has described core spliceosome members, including U1-70K (snRNP70)^22, 34, 95–97^ and disordered RBPs such as G3BP1, TIA1, and SRRM2 in tau aggregation^27, 57, 98^, co-aggregating with tau in the cytoplasm across cellular^57^, murine models^21, 47^ and in human AD brain tissue^22, 23, 96^. Most RBPs lack a strong secondary and tertiary structure and are characterized as intrinsically disordered, enabling electrostatic interactions with other proteins and diverse mRNA species to carry out their homeostatic function^99^. Notably, tau itself is highly disordered, facilitating its liquid-liquid phase separation into membrane-less organelles such as cytoplasmic stress granules or processing bodies to accelerate the fibrillization^54, 100, 101^. These TauRD interactome results further support the role of tau and associated stress granule proteins.

Current research on tau aggregation is limited to understanding additional protein interactions impacting the oligomerization and, furthermore, aggregation of tau. To address this, we sought to develop a proximity labeling system using the domain of tau detected in paired helical filaments^9, 10, 45, 102–104^. Dissecting tau interactors through immune affinity-capture is limited as it must be performed under lysis conditions preserving physiological interactions, whereas bulk cortical proteome was generated from tissues lysed in 8M urea, which can solubilize aggregated proteins. Tau interactome studies have primarily employed IP with tau antibodies^13, 15, 16, 52, 69, 105–107^, as well as capturing tau interactome by ascorbate peroxidase (APEX) proximity labeling^14, 37^. TurboID has the sensitivity to capture both transient and weakly interacting proteins, with labeling of lysine (Lys) residues in a 10 nm range within five minutes^41, 42^. TurboID is also more likely to favor labeling intrinsically disordered proteins, such as RBPs and tau itself, as the hydrophilic nature of Lys residues may allow peptides to be readily labeled^108^. This study identified several unique RBPs with an average of ∼10% K residues. Many of these TauRD interactors were found to be disease-relevant in the bulk cortex of AD and PSP across both soluble and insoluble co-aggregating proteins. Interestingly, tau IP interactomes overlapped with some insoluble-mapping RBPs across M5, M15, and M18. We hypothesize these were captured from a soluble pool of protein^22^.

sTurbo TauRD surprisingly labeled many proteins that did not overlap with the tau interactome dataset nor were reported by other tau interactomes^13, 14, 52, 69, 105, 106, 109^. Specifically, we highlight the insoluble-mapped proteins MACROH2A1, WDR82, and SF3B6, along with CHD4, SAFB, and PAPOLA, which were predominantly soluble. These findings may be influenced by several factors, including the limitations of conventional IP methods in capturing these interactions, and differences in the proximity labeling technique implemented. Overall, the sTurbo TauRD HEK293 in vitro interactome complements prior tau interactome studies outlining the physiological involvement of tau with proteasomal subunits^15, 52^, DNA-binding proteins^15^, 14-3-3 binding proteins^13, 15, 52, 69, 105^, and vesicle sorting/trafficking^14, 109^.

As a monoculture method in HEK293 cells, this study did not identify brain cell type-specific differences in tau aggregation biology. Overexpression of the TauRD may bias cells toward a stress state, especially as tau is not endogenously expressed in HEK cells and would not have native interactors seen in other commonly used neuronal lines, such as SH-SY5Y. However, MTBR inclusions seen in this model are consistent with prior studies showing stress granules or autophagosome overlap^85, 106, 110^. Further discordance of sTurbo TauRD results in comparison to other tau interactome datasets may be due to various reasons, including mutation differences (P301L, V337M, WT) influencing interactions and distinction from the tau domain (full length, 3R MTBR isoform, N-terminal region, Proline-rich domain, C-terminal region) used in this study. Lastly, we sought to build upon the previously characterized HEK-FRET TauRD biosensor aggregation cell line and extract the same amino acids of the MTBR (L243 to K375) for this model. However, further studies have demonstrated that while most of the MTBR is represented in tangles, it is not the whole fibril core^111^. Paired helical filament fibrils are composed of amino acids V336 to F378^111^. The difference in these 10 amino acids may alter this domain’s structural organization and biochemical interactions in vitro and may account for nuanced differences in interactors. Despite these clear limitations, this study contributes to our understanding of disease-relevant protein-protein interactors in Tauopathies and provides new candidate RBPs associated with tau pathology in disease^112^.

Ultimately, the sTurbo TauRD system acts as a platform for the protein aggregation field to characterize both soluble and insoluble protein-protein interaction networks related to human disease. These data also indicate that this tau proximity labeling approach provides insight into unique nuclear pathways, highlighting the importance of RBPs and spliceosome members for future mechanistic studies and potential therapeutic targets in disease.

### Data Availability Statement

The mass spectrometry proteomics RAW files and search results have been deposited to the ProteomeXchange Consortium via the PRIDE partner repository with the dataset identifier PXD059716. Supplemental Tables, raw files and FragPipe output directories from sTurbo TauRD proteomics are also available on https://www.synapse.org/Synapse:syn64396764. The 2024 human Uniprot database containing 20,597 reviewed proteins (downloaded February 7^th^, 2024, from https://www.uniprot.org/) was used for searches of mass spectrometry data. Source data are provided with this paper.

## Methods and Materials

### Experimental Design and Statistical Rationale

*Characterization of N- and CsTurbo TauRD:* Three biological replicates were collected for experiments where transfection conditions included: N-sTurbo TauRD, C-sTurbo TauRD, both N-and C-sTurbo TauRD, or reagent alone (Mock). Each transfection condition also contained a biotin treatment and biotin negative group to determine background biotinylation (i.e. endogenous biotinylated proteins).

*In vitro mass spectrometry proteomics*: Biological replicates represent different passage numbers of the same cell line, with conditions including transfected with sTurbo TauRD encoding plasmid and a mock group treated with reagent alone. Cells were prepared with 4 biological replicates in two experimental batches and a third batch consisting of 2 biological replicates, totaling 10 per transfection condition. Following quality control, 6 mock samples were analyzed for proteomics after streptavidin affinity purification. Thus, 16 cell lysates were included in the study. An *a priori* power analysis for sample size was not performed, although the highest sample size afforded was included. As previously detailed^36^, Mock peptides were run before sTurbo TauRD for mass spectrometry to reduce column contamination between sample types. Details for further statistical methods are discussed in the "*Differential abundance and ontological enrichment*” and “*Human validation analyses*” subsection.

### Cloning and plasmid isolation

Three sTurbo TauRD plasmid constructs, N-sTurbo TauRD, C-sTurbo TauRD, and P301L 2A sTurbo TauRD, were created in coordination with Emory University’s Integrated Genomics Core (EIGC). The original N- and C-terminal sTurbo fragments correspond to Addgene #153002 and Addgene #153003, respectively, with the microtubule-binding region (MTBR) based on tau amino acids 243-375 (2N4R, 441 isoform). N-sTurbo TauRD includes the MTBR, a 25 amino acid SG-rich linker, a V5 tag, N-terminal sTurbo ligase fragment, and is driven by a human Ubiquitin (hUb) promoter. C-sTurbo TauRD shares the MTBR and hUb promoter but features a 17 amino acid linker, an HA tag, and the larger C-terminal sTurbo fragment. Restriction enzyme sites were introduced with respective forward and reverse primers, outlined in **Supplemental Table S18**. The pLEX-MCS UBC promoter was a kind gift from Oskar Laur (Addgene plasmid #128045 http://n2t.net/addgene:128045;RRID:Addgene_128045). The N- and C- sTurbo TauRD PCR products were then subcloned into a FUGW backbone with a puromycin resistance ORF placed instead of GFP. The P301L 2A sTurbo TauRD PCR product was subcloned into a custom pLEX MCS vector. The P301L 2A plasmid was transformed into DH10B competent *E. coli* cells (Invitrogen, #18297010) according to manufacturer’s instructions, while DH5-a *E. coli* was used for N- and C-sTurbo TauRD plasmids. Plasmid DNA was purified following the PureLink™ HiPure Plasmid Filter Maxiprep Kit instructions (Invitrogen, #K210016) and resulting concentration measured via Nanodrop.

### Cell Maintenance

HEK293T cells (ATCC, CRL3216) were cultured in DMEM supplemented with 10% fetal bovine serum (FBS) and 1% penicillin-streptomycin at 37°C with 5% CO2, following standard protocol^36, 39, 113, 114^. For passaging, Trypsin-EDTA 0.05% (Gibco, 25300054) was utilized to detach cells, which were subsequently resuspended and washed with 1X PBS. Cell counts were performed using a hemocytometer prior to experimental plating.

### DNA Transfection

Plasmid DNA was transfected into cells at 60-80% confluency using the jetPRIME Polyplus transfection reagent (VWR, 89129) based on the manufacturer’s protocol. For 24-well dishes and 8-well chamber slides (Nunc™ Lab- Tek™ II Chamber Slide™ System, Thermo Scientific 154453), transfection was performed with 50 µL jetPRIME buffer, 1 µL jetPRIME reagent, and 0.5 µg of plasmid DNA per transfection in 500 µL of media. For co-transfection, equal amounts of N- and C-sTurbo TauRD plasmids were combined to reach the total recommended concentration. Mock transfections included the same ratio of reagents without plasmid. Cells were harvested or fixed 48 hours post-transfection.

### Biotin Treatment

On the same day as cell collection or fixation biotin labeling was performed following standard procedure, with minimal modification^39^. Biotin (Sigma, B4639) was prepared as a 10 mM stock solution in 1X PBS (pH 9.0) and stored at 4°C. A fresh 200 µM biotin solution was made by diluting the stock in serum-free (SF) media. After aspirating the complete media, cells underwent a biotin wash-out period in SF media for approximately 3 hours. Subsequently, biotin positive groups were incubated in the 200 µM biotin supplemented media for 1 hour before continuing to respective collection or fixation conditions.

### Cellular Lysis

Cells were lysed according to standard laboratory protocol^39, 113^. Briefly, cells were collected in fresh lysis buffer (8 M urea, 10 mM Tris, 100 mM NaH_2_PO_4_, pH 8.5) with 1X HALT protease/phosphatase inhibitor (Thermo Sci, 78446). Lysates were sonicated at 30% amplitude for three 5-second intervals, with the sonication tip cleaned between samples. Following centrifugation at 18,000 x g for 15 minutes at 4°C, supernatants were collected. Protein concentrations were determined using the Bicinchoninic acid assay (BCA, Pierce) and stored at -20°C.

### SDS-PAGE, Western blot (WB), and semi-quantification analysis

Lysate total protein amount was normalized and diluted to 1X Laemmli sample buffer (Biorad) and 355 mM β- mercaptoethanol (Sigma-Aldrich) as per **Supplemental Table S19**. SDS-PAGE and WB were run as previously described^39, 114, 115^. Samples were boiled and loaded onto 1.0 mm 4-12% Bis-Tris NuPage gels (NW04122, Invitrogen) alongside a molecular weight marker (New England Biosciences, P7719S or ThermoFisher, 26616). Electrophoresis was carried out in 1X MES running buffer until the molecular weight marker reached the end of the gel. Proteins were then transferred to a 0.2 µm nitrocellulose membrane at 20 V for 7 minutes using the iBlot dry transfer stack system (Invitrogen) and blocked for 30 minutes in StartingBlock Buffer (Thermo Scientific, 37543). Antibodies were diluted in StartingBlock buffer and incubated per **Supplemental Table S19.** Membranes were washed twice with TBS-T, followed by TBS washes, before imaging on the Licor Odyssey M system. Validation affinity purifications and WBs were conducted essentially as described with minor modifications. Samples with minimal non-specific binding shown by silver staining were selected for validation blots.

*WB and Silver stain intensity semi-quantification:* Protein band intensities were quantified using ImageJ FIJI software (v2.14.0, National Institutes of Health), essentially as previously described^115^.

*WB N- and C-sTurbo TauRD signals from 2A construct:* Biotinylated N- and C-sTurbo TauRD fragments, overlapping with V5 and HA signals, respectively were selected as ROIs. Intensity measurements were selected from the Streptavidin 680 blot to reduce bias from antibody signals or channel intensity variation. Inverted intensities were analyzed in GraphPad Prism (v10.1.1) the percentages of each streptavidin C-sTurbo and N- sTurbo value by the sample total were visualized as stacked bar plots.

### Immunocytochemistry (ICC) and microscopy

Acid-etched sterile coverslips (Bellco Glass SKU: 1943-10012A) were coated with Poly-L-Lysine, PLL, (Sigma P1524, MW > 300,000) according to manufacturer’s instructions. Cells were transfected with experimental plasmids, then underwent biotin labeling protocol the same day as fixation with 4% paraformaldehyde. Cells were prepared for immunofluorescence as previously reported^39^. All antibodies were diluted in antibody buffer (1X PBS in 2% normal horse serum). Antibodies for ICC were as follows: 1:500 rabbit anti-V5 (abcam: 206566), 1:500 mouse tau 2E9 (Novus Biologicals, NBP2-25162), incubated for 1 hr at room temperature and 1:300 rabbit anti-HA tag antibody (abcam: ab9110) incubated overnight at 4°C. Secondary antibodies were as follows: 1:500 Donkey anti-Rabbit FITC conjugated (Invitrogen, A16030), 1:500 Streptavidin DyLight 594 (Invitrogen, 21842), 1:500 Donkey anti-Mouse FITC conjugated (Invitrogen, A24507) and incubated at room temperature for 90 minutes in the dark. Primary and secondary only controls were incorporated to ensure specificity of fluorescence signal. All wells were counterstained with 1 µg 4′,6-diamidino-2-phenylindole, DAPI (Molecular Probes, D3571) in the first 5-minute 1X PBS wash in the dark for five minutes to stain for nuclear compartment. Cells were imaged with a 40X objective on a Keyence BZX-800 wide-field microscope.

### Streptavidin affinity purification (SA-AP)

Streptavidin affinity purification (SA-AP) was conducted according to standard protocol^35, 36, 39^ Briefly, 1 mg of protein was normalized to 500 µL with cold RIPA buffer (pH 7.5, composed of 50 mM Tris-HCl, 150 mM NaCl, 0.1% SDS, 0.5% sodium deoxycholate, and 1% Triton X-100) then underwent a 1 hour incubation at 4°C on a rotator with 83 µL pre-washed streptavidin-conjugated magnetic beads (ThermoFisher, 88817). Sequential 1 mL washes were performed to reduce non-specific protein binding to magnetic beads: RIPA for 8 minutes, 1M KCl for 8 minutes, fresh 0.1M Na_2_CO_3_ for 10s, 2M Urea in 10 mM Tris-HCl (pH 8.0) for 10s, then two 8-minute RIPA washes. Two more 1X PBS washes were performed, after which the beads were resuspended in the original 83 µL volume. Approximately 10% of the beads (8 µL) were collected, PBS removed, then resuspended in 30 µL of Laemmli sample buffer containing 2 mM biotin and 20 mM DTT. Proteins were boiled from the beads at 95°C for 15 minutes. 10 µL of this aliquot was retained for WB analysis, while 20 µL was allocated for silver stain. The remaining 75 µL beads were prepared for protease digestion and mass spectrometry.

### Silver staining procedures & semi-quantification

Silver staining was conducted before mass spectrometry and WB validation to assess the effectiveness of the streptavidin affinity purification. The silver stain protocol followed the manufacturer’s instructions (Pierce, 24612). Following fixation, washing, and staining steps, the gel was incubated in developer solution until protein ladder turned brown and protein bands were visualized the quickly rinsed with silver stain stop solution (5% acetic acid), and then incubated in fresh stop solution before imaging on the Odyssey M with the RGB Trans channel.

*Silver stain quality control gel:* FIJI intensities were obtained from the entire lane as ROI. These values were assessed in GraphPad Prism and visualized in a scatter dot plot for Mock and sTurbo TauRD groups. Mock intensities above background levels were excluded from further analysis.

### Protease Digestion

Mock transfection and sTurbo TauRD HEK293T SA-AP samples were digested essentially as described^35, 36, 39^. Briefly, samples were normalized to 300 µL in 100 mM Tris-HCl pH 8 buffer. Lysates were reduced with 1 mM dithiothreitol (DTT) and alkylated with 5 mM iodoacetamide (IAA) at room temperature for 30 minutes, respectively. Proteins were digested overnight at room temperature with 2 µg Lys-C protease (Wako). The next day, 2 µg Trypsin was added and incubated overnight. The following day, enzyme activity was quenched with acidifying buffer (1:9, v/v; 10% formic acid and 1% trifluoroacetic acid) and desalted by loading the peptides onto a 10 mg Oasis PRiME HLB 96-well plate (Waters), followed by washing twice with Buffer A (0.1% TFA) and then eluted with Buffer C (50% acetonitrile, ACN and 0.1% TFA). Desalted peptides were lyophilized with a CentriVap Centrifugal Vacuum Concentrator (Labconco) overnight.

### Liquid Chromatography – tandem mass spectrometry (LC-MS/MS)

Peptides were resuspended and sonicated in Buffer A for loading. 20 uL was loaded onto an Evosep One tip, then separated by an Evosep One Liquid Chromatography (LC) system using a 15 cm × 150 μm i.d. Water’s 1.7 µm CSH capillary with a preprogrammed 88 min gradient procedure (15 samples per day). MS procedure was executed essentially as described^35, 36^. MS was performed with an Orbitrap Fusion Lumos Tribrid (ThermoFisher) in positive ion mode for data dependent acquisition (DDA) MS. The mass spectrometer was run at top speed mode with each cycle lasting for 3 seconds and consisted of one full MS scan and as many MS/MS events as possible within the allowed cycle time. MS scans were collected at a resolution of 60,000, with a 400 to 1600 m/z range, 4 × 10^5^ automatic gain control (AGC), and a maximum ion injection time of 118 ms. All higher energy collision-induced dissociation (HCD) MS/MS spectra were acquired at a resolution of 30,000, with a 1.6 m/z isolation width, 30% collision energy, 5 × 10^4^ AGC target, and 54 ms maximum ion injection time. Previously sequenced peaks were dynamically excluded for 30 s within a 10-ppm isolation window.

### Database search and quantification

Raw files were searched with FragPipe (v 21.1). FragPipe relies on its integration with MSFragger (v 4.0) for peptide identification from a closed database. The proteins were identified by searching against the February 2024 canonical Uniprot database with 20,597 proteins and 4 independent protein sequences aligning to the N- sTurbo, C-sTurbo, V5 tag, and HA tag ORFs to identify the sTurbo TauRD recombinant protein. 51 total contaminant sequences and all 20,652 reverse sequences were added for false discovery rate (FDR) calculation, which was set to 1%. The default FragPipe Label-Free Quantitation – Match Between Runs (LFQ-MBR) workflow was loaded with default parameters with minimal modifications.

Percolator (v3.6.4) filtered peptide spectral matches (PSMs) using a support vector machine algorithm to control PSMs matched to peptides from decoy proteins and best discern true PSMs to be kept in final protein assembly. Peptide-to-spectra matches were rescored with MSBooster and Percolator for predicting retention time and spectra with a minimum probability of 0.5. Briefly, precursor mass tolerance was -20 to 20 ppm, fragment mass tolerance of 20 ppm, with mass calibration and parameter optimization selected, and isotope error was set to 0/1/2. Enzyme specificity was set to strict-trypsin with up to 4 missed cleavages allowed due to the potential lysine-biotin modification impacting tryptic cleavage. Peptide length ranged from 7 to 50 amino acids, and peptide mass from 500 to 5,000 Da. Variable modifications included: methionine oxidation (+15.9949 Da), N-terminal acetylation (+42.0106 Da), and phosphorylation on STY residues (+79.96633 Da) with a maximum of 3 variable modifications per peptide. Fixed modification included cysteine carbamidomethylation (+57.02146 Da).

Quantification was conducted by the IonQuant module (v1.10.12) with MBR permitted. Abundances were also reported using with the topN method with topN peaks set to 200. IonQuant parameters were as follows: 2 minimum ions, 3 minimum scans, minimum matched fragments of 4, and a retention time tolerance of 0.4 min. After FDR filtering with Philosopher (v 5.1.0), a total of 60,847 peptide spectral matches (PSMs), 54,148 modified peptides, 50,674 peptides, and 5,262 protein groups were detected.

### Differential abundance and ontological enrichment

After filtering out proteins absent in 50% or more across samples^18^, 2,167 proteins were retained for downstream analyses. An in-house function performed sample-wise imputation of missing values similar to the Perseus program^116^ in R (v 4.3.3, 2024-02-29). Briefly, values were imputed according to a normal distribution falling between +/- 0.3 standard deviations from the noise level. The noise level was assumed to be -1.8 standard deviations from the mean of the Log_2_(LFQ intensity) values. The resulting abundance matrix was used for differential abundance and bioinformatics analyses. Raw intensity values are provided for pre- and post- missingness thresholding (**Supplemental Tables S1** & **S2**). A Student’s two-tailed t-test was implemented to determine differentially abundant (DEA) proteins across groups (**Supplemental Table S6**). Volcano plots of - log_10_ unadjusted p-values were generated using the custom plotVolc function, accessible via https://www.github.com/edammer/parANOV<u>^A^</u>. This script also provides the parANOVA.dex function for fast parallel calculation of one-way analysis of variance (ANOVA) with Tukey-corrected pairwise comparison p values when there are more than 2 groups compared. Bonferroni correction was employed instead of Tukey post hoc test in cases where Tukey p value estimates were less than 1x10^-8.5^, avoiding a ceiling effect due to imprecise small Tukey p value estimation.

Ontologies enriched by module were determined by Fisher’s exact test (FET) of enriched sTurbo TauRD proteins using the GOparallel function. This script calculates one-tailed enrichment p value for input lists into over 13,000 human GO annotated gene lists from a GMT-formatted file obtained from the Bader Lab site December 1^st^ 2023, as previously described^76, 115, 117–124^. Supporting online resources are available via https://www.github.com/edammer/GOparallel.

### Human Brain tissue proteomics

#### University of Pennsylvania (UPenn) Protein Quantification and Subset by Disease Condition

The University of Pennsylvania cohort raw files and metadata were obtained from synapse.org syn53177242. Brain tissue from the dorsolateral prefrontal cortex (BA 9) preparation, isobaric Tandem Mass Tag (TMT) labeling, subsequent mass spectrometry data acquisition, and protein quantification is originally detailed in Shantaraman et al, 2024^125^. All 364 UPenn samples underwent technical artifact batch normalization using a Tunable Approach for Median Polish of Ratio (TAMPOR)^126^ leveraging the central tendency of only true GIS samples for achieving the final converged log2(abundance) central tendency at 0 relative log2(abundance) units. Then, the resulting abundance matrix was subset to include control (n=46), Alzheimer’s Disease (n=49), and Progressive Supranuclear Palsy (n=26) cases. Age, sex, and postmortem interval (PMI) covariates were then modeled and covariance removed via non-parametric bootstrap regression with the protein-specific coefficient for each modelled variable set to the median from 1000 bootstraps^127^.

#### Weighted Gene Co-expression Network Analysis (WGCNA)

The WGCNA algorithm^79^ was applied to execute co-expression network analysis to obtain modules of highly correlated protein groups of cases with neuropathological diagnoses of AD, PSP, and control, as described^18, 19, 56, 76, 77, 125^. The WGCNA::blockwiseModules() function was implemented for network generation. The following settings were used: soft threshold power = 9, deepSplit = 2, minimum module size = 25, merge cut height = 0.07, mean topological overlap matrix (TOM) denominator, signed network with partitioning about medioids (PAM) per the dendrogram, and a module reassignment threshold of p < 0.05. The signedkME function was also employed to determine biweight midcorrelation (bicor) of protein to module membership (**Supplemental Table S11**). Proteins with high kME values are considered to be module hubs. Module i-Graphs were generated with the buildIgraphs function, available from https://www.github.com/edammer/netOps. i-Graphs with protein-protein interaction database (BioGrid) integration contain information for sTurbo TauRD interacting proteins across network hubs **(Supplementary** Fig. 4-5). Additional visualization layout and formatting for dataset hits were performed using Illustrator 2024.

#### Fisher’s Exact Test (FET) analyses

*Gene Ontology (GO):* A hypergeometric overlap test was performed for each module protein’s membership using the above-detailed GOparallel function and human reference GMT file.

*Cell Type Marker Enrichment*: One-tailed Fisher’s exact test FDR adjusted -log_10_(p-values) for the enrichment of brain cell type marker proteins in human AD & PSP bulk frontal cortex proteomics modules were visualized as heatmaps using the geneListFET function from open source code available online at https://www.github.com/edammer/CellTypeFET, as previously published^17, 119, 121, 123–125, 128–133^

*Cellular Compartment Enrichment Analyses:* This geneListFET script was adapted to visualize the enrichment of module proteins to a curated list of subcellular components downloaded from Uniprot November 2024 (Subcellular localization GO-SL IDs: 0516, 0515, 0243, 0090, 0091, 0191, 0048, 0158, 0132, 0173, 0039) and enrichment of sarkosyl-insoluble TMT-labeled proteins from AD and CTL postmortem brain^84^.

*Tau interactome mapping to Human network:* AD enriched tau interactomes (Tau-5 and PHF-1 epitope-recognizing antibody immunoenriched proteomes) from human postmortem brain tissue^15, 16^ and *in vitro* sTurbo TauRD interactors to human brain proteomic modules were separately considered as input lists for independent runs of the geneListFET against module membership.

*Insoluble TMT proteomics to Human network:* A previously published dataset of sarkosyl insoluble TMT proteomics from AD and CTL brains were mapped to the human network modules to define “insoluble mapping” proteins. Enrichment was visualized as a barplot of the -log_10_(unadjusted p-values). Insoluble mapping proteins were manually annotated within module-specific i-Graph(s) in Illustrator.

#### RNA-binding protein (RBP) module analyses and visualization

sTurbo TauRD mapping RBP-enriched module (M5, M15, and M18) gene IDs were merged with a comprehensive RBP list^134^ accessed via https://apps.embl.de/rbpbase/ and downloaded June 2024. For visualizing TauRD interactors vs Mock transfected controls, LFQ intensities were normalized by z-score and scaled to transform the data ranges from -4 to +4. The resulting matrix was visualized in R using the ComplexHeatmap library. To annotate RBP types associated with RBP-enriched modules, TauRD interactome was merged with previously published data describing 356 annotated RBPs for respective function, localization, and amino acid motifs^86^ and visualized via ggplot().

## Author Contributions

Conceptualization (N.T.S., S.R., S.M.S.); Methodology (N.T.S., S.R., S.M.S., C.A.B.); Software (S.M.S., A.S., E.B.D.); Validation (S.M.S., A.S., M.A.K.); Formal analysis (S.M.S., A.S.); Investigation (S.M.S., M.A.K., L.Z., D.M.D., C.A.B., P.B., Q.G.); Resources (S.R., N.T.S.); Data Curation (S.M.S., A.S., E.B.D., D.M.D.); Writing: original draft (S.M.S., N.T.S.); Writing: review & editing (S.M.S.); Visualization (S.M.S); Supervision (N.T.S., S.R); Project administration (S.M.S., N.T.S., D.M.D); Funding acquisition (S.M.S., N.T.S.).

## Acknowledgements

This study was supported in part by the Emory Integrated Genomics Core (EIGC) (RRID:SCR_023529), which is subsidized by the Emory University School of Medicine and is one of the Emory Integrated Core Facilities. Additional support was provided by the Georgia Clinical & Translational Science Alliance of the National Institutes of Health under Award Number UL1TR002378. The content is solely the responsibility of the authors and does not necessarily reflect the official views of the National Institutes of Health.

Financial support for this research was provided by NIH NIA grants U01AG061357 (N.T.S.), R01AG075820 (N.T.S. & S.R.), and F31AG079670 (S.M.S.). Research was supported in part by the Emory Integrated Proteomics Core (EIPC), Integrated Cellular Imaging (ICI) Core, and the Emory Integrated Genomics Core (EIGC) shared resources of Winship Cancer Institute of Emory University and NIH/NCI under award number P30CA138292. Specifically, I would like to acknowledge the support of Laura Fox & Gaurav Joshi (ICI), and Oskar Laur (EIGC) for their expertise & contributions to this project.

---

**Supplemental Figure S1.**
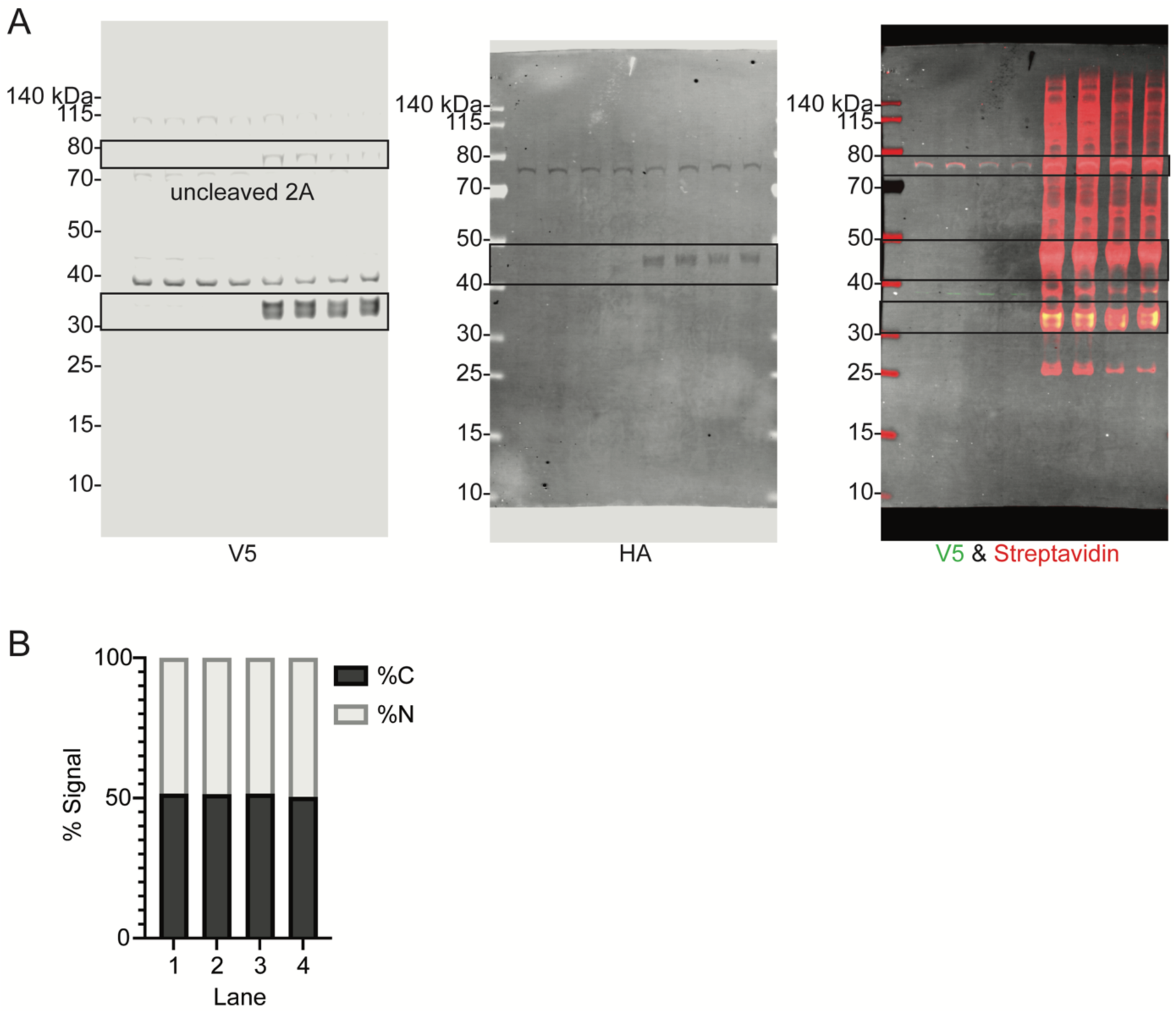
2A efficiently cleaves fragments and equally biotinylates sTurbo TauRD fragments. **(A)** Cells were transfected with either reagent alone or with sTurbo P301L TauRD 2A plasmid DNA and underwent standard biotin labeling before 8M Urea lysis and protein collection. WB analysis identified recombinant proteins (V5 and HA) and biotinylated proteins. sTurbo TauRD cleavage products were observed at expected molecular weights: ∼27 kDa for N- sTurbo TauRD (V5) and ∼43 kDa for C-sTurbo TauRD (HA). A faint, uncleaved product (∼73 kDa) was observed on the V5 blot. The streptavidin blot also displayed self-biotinylation of the cleaved protein products (HA, white; V5, green). **(B)** Further characterization of the 2A recombinant protein ligase activity reveals the N- and C-sTurbo biotinylated fragments are equally abundant at each ∼50% signal via western blot analysis of streptavidin overlaying biochemical tag protein band.

**Supplemental Figure S2.**
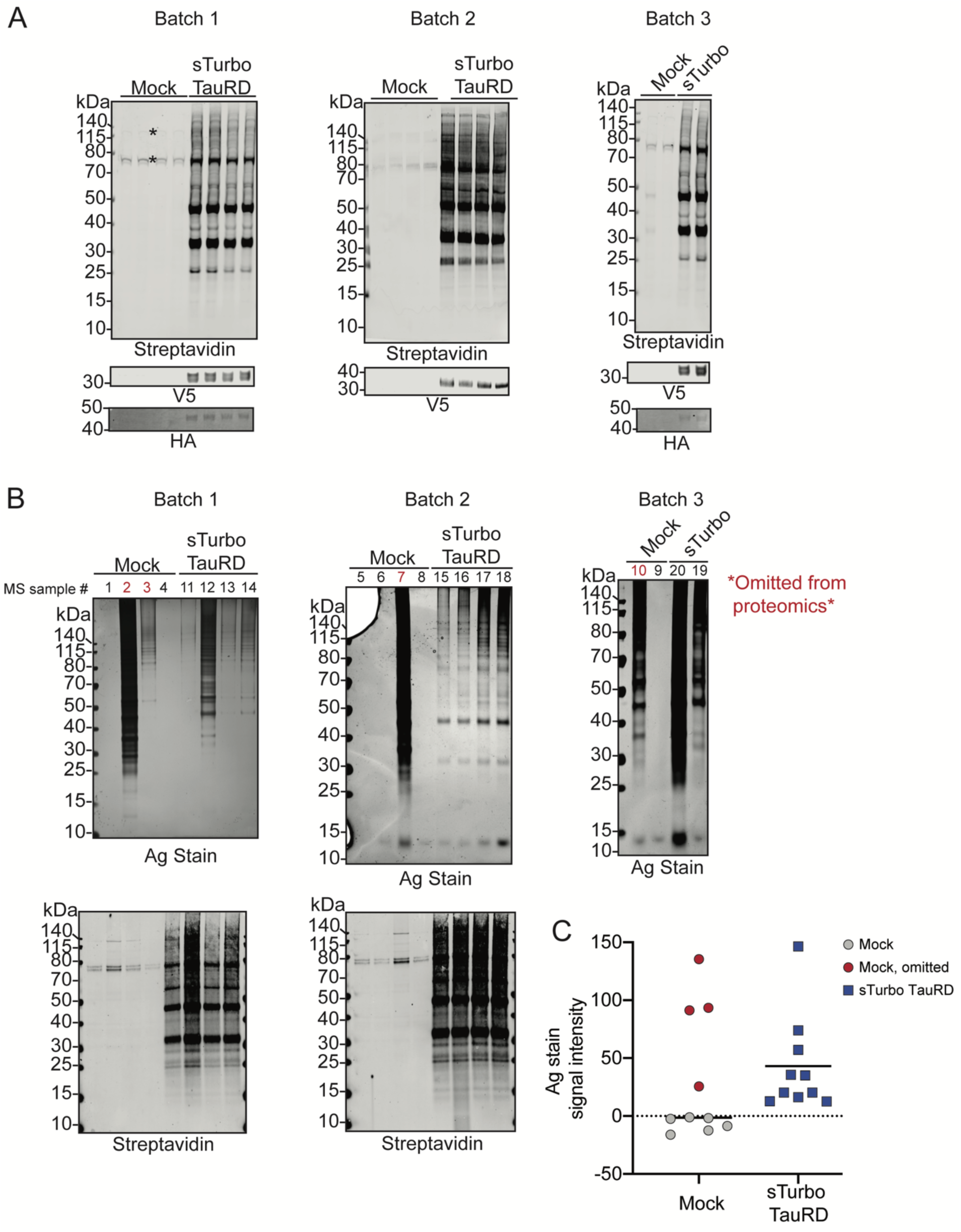
Quality control western blots and total protein silver stains. **(A)** Total lysate input (20 µg) of sTurbo TauRD from HEK293T cells shows robust and consistent labeling across replicates and batches, shown by streptavidin blots. V5 and HA recombinant protein tags were also probed to confirm expression of sTurbo TauRD across lysates. **(B)** Silver stain (Ag stain) displaying total protein after streptavidin affinity purification shows varying total signal in sTurbo TauRD samples and has enriched biotinylated proteins, shown through streptavidin blot. Mock samples with non-specific binding to streptavidin beads, and resulting positive signal found in silver stain, were omitted from this proteomics study (red). Signal intensities from the Ag stain gel lanes are depicted in panel **(C)**.

**Supplemental Figure S3.**
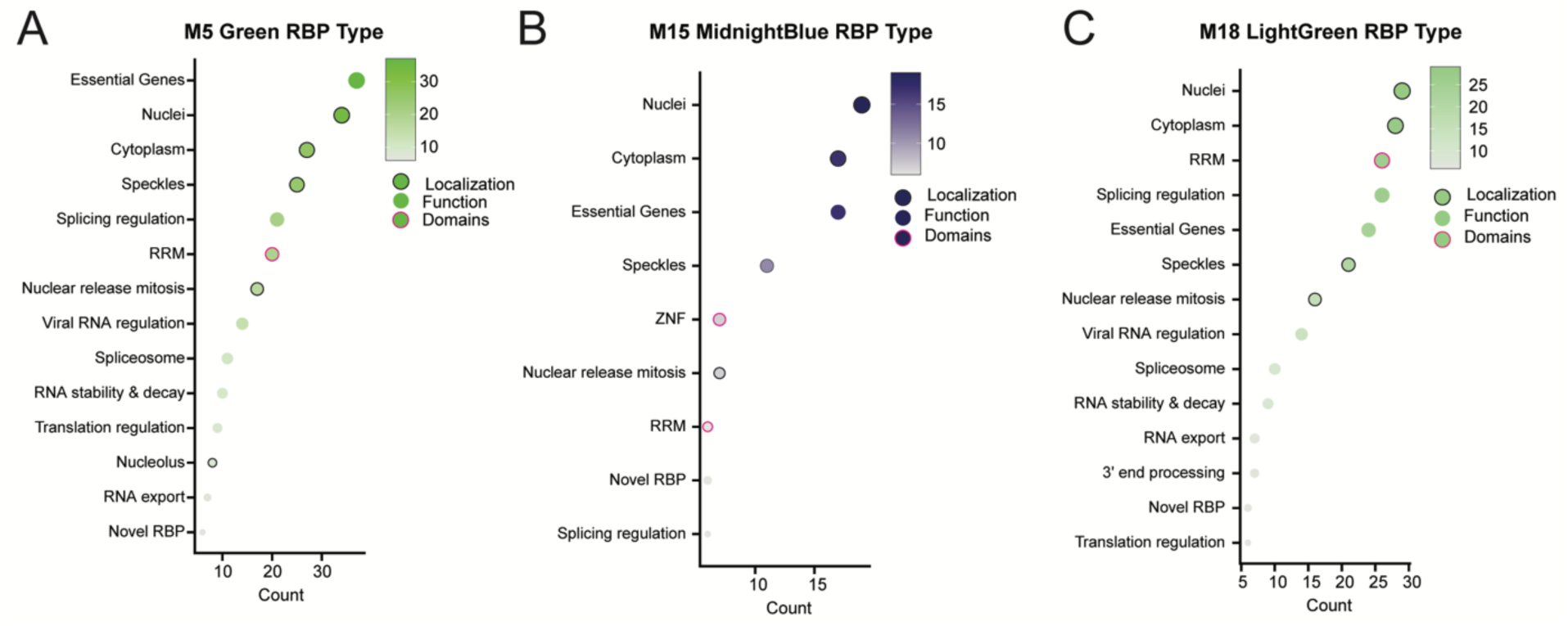
Insoluble disease mapping module classification across RBP localization, key functions, and domains. Module members across RBP-enriched modules M5 **(A)**, M15 **(B)**, and M18 **(C)** were integrated with a list of 356 well-characterized RBPs to delineate common and divergent features across each module. Count of module RBPs mapping to each characteristic is visualized.

**Supplemental Figure S4.**
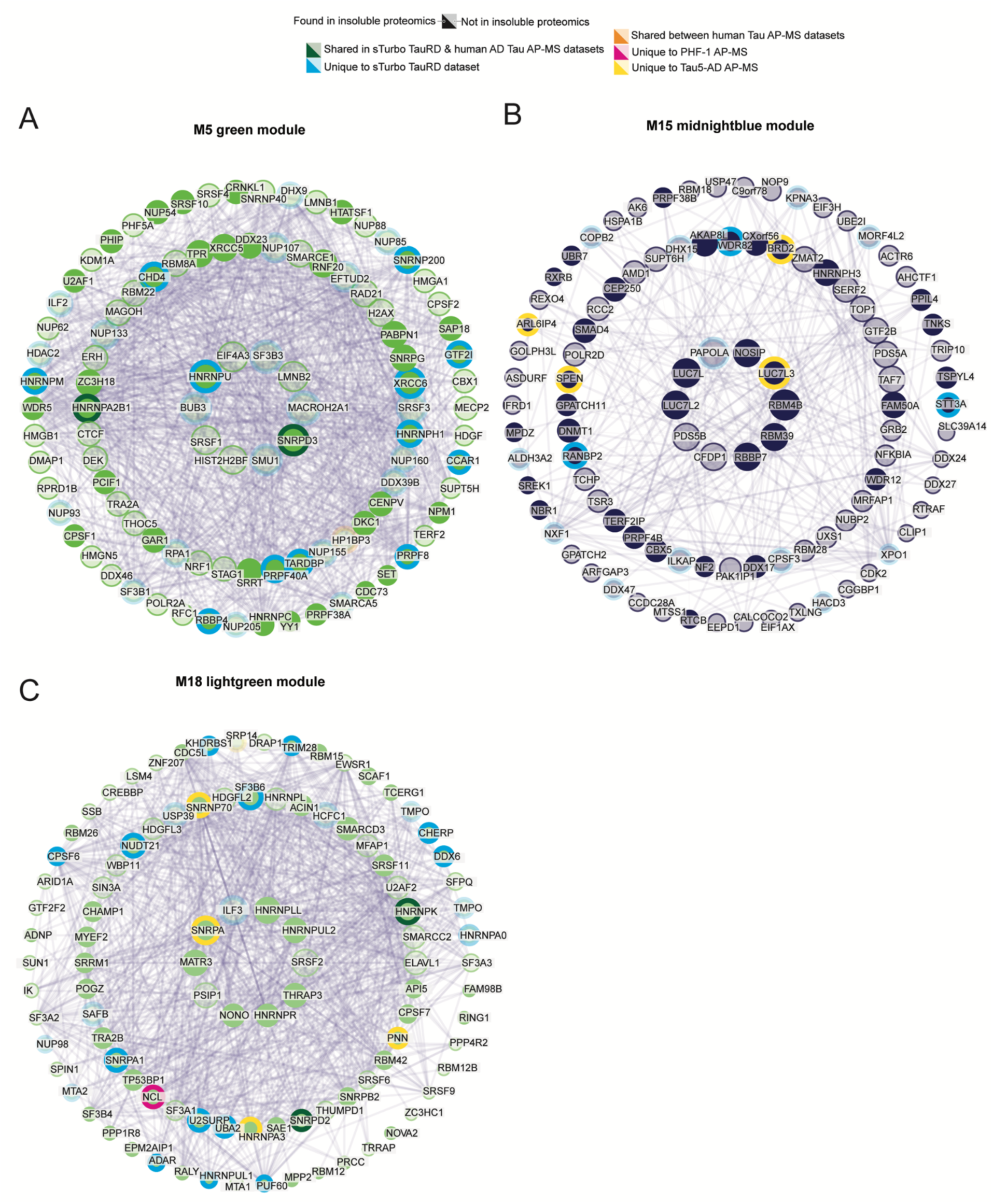
Insoluble and RBP-enriched module hub proteins and BioGrid interaction networks. Biogrid integration in which edge lines provide information on biological interactions across hub proteins for insoluble disease-related modules, M18 **(A)**, M5 **(B)** and M15 **(C).** Protein nodes are colored if Tau interactome proteins are identified within the module, either from sTurbo TauRD (unique, blue; shared with AD Tau AP-MS, green) or only described in human Tau AD AP-MS interactomes (warm colors). sTurbo TauRD uniquely captures hub proteins across all three modules, including ILF3, MACROH2A1, SF3B6, and PAPOLA. Shared proteins across Tau interactome datasets include RBPs HNRNPK, SNRPD2, HNRNPA2B1.

## References

(1) Neve, R. L.; Harris, P.; Kosik, K. S.; Kurnit, D. M.; Donlon, T. A. Identification of cDNA clones for the human microtubule-associated protein tau and chromosomal localization of the genes for tau and microtubule-associated protein 2. Brain Res 1986, 387 (3), 271–280. DOI: 10.1016/0169-328x(86)90033-1 From NLM Medline.

(2) Goedert, M.; Spillantini, M. G.; Potier, M. C.; Ulrich, J.; Crowther, R. A. Cloning and sequencing of the cDNA encoding an isoform of microtubule-associated protein tau containing four tandem repeats: differential expression of tau protein mRNAs in human brain. EMBO J 1989, 8 (2), 393–399. DOI: 10.1002/j.1460-2075.1989.tb03390.x.

(3) Panda, D.; Goode, B. L.; Feinstein, S. C.; Wilson, L. Kinetic stabilization of microtubule dynamics at steady state by tau and microtubule-binding domains of tau. Biochemistry 1995, 34 (35), 11117–11127. DOI: 10.1021/bi00035a017 From NLM Medline.

(4) Drechsel, D. N.; Hyman, A. A.; Cobb, M. H.; Kirschner, M. W. Modulation of the dynamic instability of tubulin assembly by the microtubule-associated protein tau. Mol Biol Cell 1992, 3 (10), 1141–1154. DOI: 10.1091/mbc.3.10.1141 From NLM Medline.

(5) Spillantini, M. G.; Goedert, M. Tau protein pathology in neurodegenerative diseases. Trends Neurosci 1998, 21 (10), 428–433. DOI: 10.1016/s0166-2236(98)01337-x From NLM Medline.

(6) Arendt, T.; Stieler, J. T.; Holzer, M. Tau and tauopathies. Brain Res Bull 2016, 126 (Pt 3), 238–292. DOI: 10.1016/j.brainresbull.2016.08.018 From NLM Medline.

(7) Spillantini, M. G.; Murrell, J. R.; Goedert, M.; Farlow, M. R.; Klug, A.; Ghetti, B. Mutation in the tau gene in familial multiple system tauopathy with presenile dementia. Proceedings of the National Academy of Sciences 1998, 95 (13), 7737–7741. DOI: 10.1073/pnas.95.13.7737.

(8) Hutton, M.; Lendon, C. L.; Rizzu, P.; Baker, M.; Froelich, S.; Houlden, H.; Pickering-Brown, S.; Chakraverty, S.; Isaacs, A.; Grover, A.;, et al. Association of missense and 5’-splice-site mutations in tau with the inherited dementia FTDP-17. Nature 1998, 393 (6686), 702–705. DOI: 10.1038/31508 From NLM Medline.

(9) Nukina, N.; Ihara, Y. One of the antigenic determinants of paired helical filaments is related to tau protein. J Biochem 1986, 99 (5), 1541–1544. DOI: 10.1093/oxfordjournals.jbchem.a135625 From NLM Medline.

(10) Kosik, K. S.; Joachim, C. L.; Selkoe, D. J. Microtubule-associated protein tau (tau) is a major antigenic component of paired helical filaments in Alzheimer disease. Proc Natl Acad Sci U S A 1986, 83 (11), 4044–4048. DOI: 10.1073/pnas.83.11.4044 From NLM Medline.

(11) Serrano-Pozo, A.; Frosch, M. P.; Masliah, E.; Hyman, B. T. Neuropathological alterations in Alzheimer disease. Cold Spring Harb Perspect Med 2011, 1 (1), a006189. DOI: 10.1101/cshperspect.a006189 From NLM Medline.

(12) Askenazi, M.; Kavanagh, T.; Pires, G.; Ueberheide, B.; Wisniewski, T.; Drummond, E. Compilation of reported protein changes in the brain in Alzheimer’s disease. Nat Commun 2023, 14 (1), 4466. DOI: 10.1038/s41467-023-40208-x From NLM Medline.

(13) Betters, R. K.; Luhmann, E.; Gottschalk, A. C.; Xu, Z.; Shin, M. R.; Ptak, C. P.; Fiock, K. L.; Radoshevich, L. C.; Hefti, M. M. Characterization of the Tau Interactome in Human Brain Reveals Isoform-Dependent Interaction with 14-3-3 Family Proteins. eNeuro 2023, 10 (3). DOI: 10.1523/ENEURO.0503-22.2023 From NLM Medline.

(14) Tracy, T. E.; Madero-Perez, J.; Swaney, D. L.; Chang, T. S.; Moritz, M.; Konrad, C.; Ward, M. E.; Stevenson, E.; Huttenhain, R.; Kauwe, G.;, et al. Tau interactome maps synaptic and mitochondrial processes associated with neurodegeneration. Cell 2022, 185 (4), 712–728 e714. DOI: 10.1016/j.cell.2021.12.041 From NLM Medline.

(15) Drummond, E.; Pires, G.; MacMurray, C.; Askenazi, M.; Nayak, S.; Bourdon, M.; Safar, J.; Ueberheide, B.; Wisniewski, T. Phosphorylated tau interactome in the human Alzheimer’s disease brain. Brain 2020, 143 (9), 2803–2817. DOI: 10.1093/brain/awaa223 From NLM Medline.

(16) Hsieh, Y. C.; Guo, C.; Yalamanchili, H. K.; Abreha, M.; Al-Ouran, R.; Li, Y.; Dammer, E. B.; Lah, J. J.; Levey, A. I.; Bennett, D. A.;, et al. Tau-Mediated Disruption of the Spliceosome Triggers Cryptic RNA Splicing and Neurodegeneration in Alzheimer’s Disease. Cell Rep 2019, 29 (2), 301–316 e310. DOI: 10.1016/j.celrep.2019.08.104 From NLM Medline.

(17) Johnson, E. C. B.; Dammer, E. B.; Duong, D. M.; Yin, L.; Thambisetty, M.; Troncoso, J. C.; Lah, J. J.; Levey, A. I.; Seyfried, N. T. Deep proteomic network analysis of Alzheimer’s disease brain reveals alterations in RNA binding proteins and RNA splicing associated with disease. Mol Neurodegener 2018, 13 (1), 52. DOI: 10.1186/s13024-018-0282-4 From NLM Medline.

(18) Johnson, E. C. B.; Carter, E. K.; Dammer, E. B.; Duong, D. M.; Gerasimov, E. S.; Liu, Y.; Liu, J.; Betarbet, R.; Ping, L.; Yin, L.;, et al. Large-scale deep multi-layer analysis of Alzheimer’s disease brain reveals strong proteomic disease-related changes not observed at the RNA level. Nat Neurosci 2022, 25 (2), 213–225. DOI: 10.1038/s41593-021-00999-y From NLM Medline.

(19) Johnson, E. C. B.; Dammer, E. B.; Duong, D. M.; Ping, L.; Zhou, M.; Yin, L.; Higginbotham, L. A.; Guajardo, A.; White, B.; Troncoso, J. C.;, et al. Large-scale proteomic analysis of Alzheimer’s disease brain and cerebrospinal fluid reveals early changes in energy metabolism associated with microglia and astrocyte activation. Nat Med 2020, 26 (5), 769–780. DOI: 10.1038/s41591-020-0815-6.

(20) Lester, E.; Parker, R. Tau, RNA, and RNA-Binding Proteins: Complex Interactions in Health and Neurodegenerative Diseases. Neuroscientist 2023, 10738584231154551. DOI: 10.1177/10738584231154551 From NLM.

(21) Maziuk, B. F.; Apicco, D. J.; Cruz, A. L.; Jiang, L.; Ash, P. E. A.; da Rocha, E. L.; Zhang, C.; Yu, W. H.; Leszyk, J.; Abisambra, J. F.;, et al. RNA binding proteins co-localize with small tau inclusions in tauopathy. Acta Neuropathol Commun 2018, 6 (1), 71. DOI: 10.1186/s40478-018-0574-5 From NLM Medline.

(22) Bai, B.; Hales, C. M.; Chen, P. C.; Gozal, Y.; Dammer, E. B.; Fritz, J. J.; Wang, X.; Xia, Q.; Duong, D. M.; Street, C.;, et al. U1 small nuclear ribonucleoprotein complex and RNA splicing alterations in Alzheimer’s disease. Proc Natl Acad Sci U S A 2013, 110 (41), 16562–16567. DOI: 10.1073/pnas.1310249110 From NLM Medline.

(23) Diner, I.; Hales, C. M.; Bishof, I.; Rabenold, L.; Duong, D. M.; Yi, H.; Laur, O.; Gearing, M.; Troncoso, J.; Thambisetty, M.;, et al. Aggregation properties of the small nuclear ribonucleoprotein U1-70K in Alzheimer disease. J Biol Chem 2014, 289 (51), 35296–35313. DOI: 10.1074/jbc.M114.562959 From NLM Medline.

(24) Hsieh, Y. C.; Guo, C.; Yalamanchili, H. K.; Abreha, M.; Al-Ouran, R.; Li, Y.; Dammer, E. B.; Lah, J. J.; Levey, A. I.; Bennett, D. A.;, et al. Tau-Mediated Disruption of the Spliceosome Triggers Cryptic RNA Splicing and Neurodegeneration in Alzheimer’s Disease. Cell Rep 2019, 29 (2), 301–316.e310. DOI: 10.1016/j.celrep.2019.08.104 From NLM.

(25) Kavanagh, T.; Halder, A.; Drummond, E. Tau interactome and RNA binding proteins in neurodegenerative diseases. Mol Neurodegener 2022, 17 (1), 66. DOI: 10.1186/s13024-022-00572-6 From NLM Medline.

(26) Lester, E.; Ooi, F. K.; Bakkar, N.; Ayers, J.; Woerman, A. L.; Wheeler, J.; Bowser, R.; Carlson, G. A.; Prusiner, S. B.; Parker, R. Tau aggregates are RNA-protein assemblies that mislocalize multiple nuclear speckle components. Neuron 2021, 109 (10), 1675–1691.e1679. DOI: 10.1016/j.neuron.2021.03.026 From NLM.

(27) Ash, P. E. A.; Lei, S.; Shattuck, J.; Boudeau, S.; Carlomagno, Y.; Medalla, M.; Mashimo, B. L.; Socorro, G.; Al-Mohanna, L. F. A.; Jiang, L.;, et al. TIA1 potentiates tau phase separation and promotes generation of toxic oligomeric tau. Proc Natl Acad Sci U S A 2021, 118 (9). DOI: 10.1073/pnas.2014188118 From NLM Medline.

(28) LeBlang, C. J.; Medalla, M.; Nicoletti, N. W.; Hays, E. C.; Zhao, J.; Shattuck, J.; Cruz, A. L.; Wolozin, B.; Luebke, J. I. Reduction of the RNA Binding Protein TIA1 Exacerbates Neuroinflammation in Tauopathy. Front Neurosci 2020, 14, 285. DOI: 10.3389/fnins.2020.00285 From NLM PubMed-not-MEDLINE.

(29) Apicco, D. J.; Ash, P. E. A.; Maziuk, B.; LeBlang, C.; Medalla, M.; Al Abdullatif, A.; Ferragud, A.; Botelho, E.; Ballance, H. I.; Dhawan, U.;, et al. Reducing the RNA binding protein TIA1 protects against tau-mediated neurodegeneration in vivo. Nat Neurosci 2018, 21 (1), 72–80. DOI: 10.1038/s41593-017-0022-z From NLM Medline.

(30) Vanderweyde, T.; Apicco, D. J.; Youmans-Kidder, K.; Ash, P. E. A.; Cook, C.; Lummertz da Rocha, E.; Jansen-West, K.; Frame, A. A.; Citro, A.; Leszyk, J. D.;, et al. Interaction of tau with the RNA-Binding Protein TIA1 Regulates tau Pathophysiology and Toxicity. Cell Rep 2016, 15 (7), 1455–1466. DOI: 10.1016/j.celrep.2016.04.045 From NLM Medline.

(31) Jiang, L.; Lin, W.; Zhang, C.; Ash, P. E. A.; Verma, M.; Kwan, J.; van Vliet, E.; Yang, Z.; Cruz, A. L.; Boudeau, S.;, et al. Interaction of tau with HNRNPA2B1 and N(6)-methyladenosine RNA mediates the progression of tauopathy. Mol Cell 2021, 81 (20), 4209–4227.e4212. DOI: 10.1016/j.molcel.2021.07.038 From NLM.

(32) Montalbano, M.; McAllen, S.; Puangmalai, N.; Sengupta, U.; Bhatt, N.; Johnson, O. D.; Kharas, M. G.; Kayed, R. RNA-binding proteins Musashi and tau soluble aggregates initiate nuclear dysfunction. Nat Commun 2020, 11 (1), 4305. DOI: 10.1038/s41467-020-18022-6 From NLM Medline.

(33) Younas, N.; Zafar, S.; Shafiq, M.; Noor, A.; Siegert, A.; Arora, A. S.; Galkin, A.; Zafar, A.; Schmitz, M.; Stadelmann, C.;, et al. SFPQ and Tau: critical factors contributing to rapid progression of Alzheimer’s disease. Acta Neuropathol 2020, 140 (3), 317–339. DOI: 10.1007/s00401-020-02178-y From NLM.

(34) Bishof, I.; Dammer, E. B.; Duong, D. M.; Kundinger, S. R.; Gearing, M.; Lah, J. J.; Levey, A. I.; Seyfried, N. T. RNA-binding proteins with basic-acidic dipeptide (BAD) domains self-assemble and aggregate in Alzheimer’s disease. J Biol Chem 2018, 293 (28), 11047–11066. DOI: 10.1074/jbc.RA118.001747 From NLM Medline.

(35) Rayaprolu, S.; Bitarafan, S.; Santiago, J. V.; Betarbet, R.; Sunna, S.; Cheng, L.; Xiao, H.; Nelson, R. S.; Kumar, P.; Bagchi, P.;, et al. Cell type-specific biotin labeling in vivo resolves regional neuronal and astrocyte proteomic differences in mouse brain. Nat Commun 2022, 13 (1), 2927. DOI: 10.1038/s41467-022-30623-x From NLM Medline.

(36) Sunna, S.; Bowen, C.; Zeng, H.; Rayaprolu, S.; Kumar, P.; Bagchi, P.; Dammer, E. B.; Guo, Q.; Duong, D. M.; Bitarafan, S.;, et al. Cellular Proteomic Profiling Using Proximity Labeling by TurboID-NES in Microglial and Neuronal Cell Lines. Mol Cell Proteomics 2023, 22 (6), 100546. DOI: 10.1016/j.mcpro.2023.100546 From NLM Medline.

(37) Batra, S.; Vaquer-Alicea, J.; Valdez, C.; Taylor, S. P.; Manon, V. A.; Vega, A. R.; Kashmer, O. M.; Kolay, S.; Lemoff, A.; Cairns, N. J.;, et al. VCP regulates early tau seed amplification via specific cofactors. Mol Neurodegener 2025, 20 (1), 2. DOI: 10.1186/s13024-024-00783-z From NLM Medline.

(38) DeCaprio, J.; Kohl, T. O. Immunoprecipitation. Cold Spring Harb Protoc 2020, 2020 (11). DOI: 10.1101/pdb.top098509 From NLM Medline.

(39) Bowen, C. A.; Nguyen, H. M.; Lin, Y.; Bagchi, P.; Natu, A.; Espinosa-Garcia, C.; Werner, E.; Kumari, R.; Brandelli, A. D.; Kumar, P.;, et al. Proximity Labeling Proteomics Reveals Kv1.3 Potassium Channel Immune Interactors in Microglia. Mol Cell Proteomics 2024, 23 (8), 100809. DOI: 10.1016/j.mcpro.2024.100809 From NLM Medline.

(40) Qin, W.; Cho, K. F.; Cavanagh, P. E.; Ting, A. Y. Deciphering molecular interactions by proximity labeling. Nat Methods 2021, 18 (2), 133–143. DOI: 10.1038/s41592-020-01010-5 From NLM Medline.

(41) Cho, K. F.; Branon, T. C.; Rajeev, S.; Svinkina, T.; Udeshi, N. D.; Thoudam, T.; Kwak, C.; Rhee, H. W.; Lee, I. K.; Carr, S. A.;, et al. Split-TurboID enables contact-dependent proximity labeling in cells. Proc Natl Acad Sci U S A 2020, 117 (22), 12143–12154. DOI: 10.1073/pnas.1919528117 From NLM Medline.

(42) Cho, K. F.; Branon, T. C.; Udeshi, N. D.; Myers, S. A.; Carr, S. A.; Ting, A. Y. Proximity labeling in mammalian cells with TurboID and split-TurboID. Nat Protoc 2020, 15 (12), 3971–3999. DOI: 10.1038/s41596-020-0399-0 From NLM Medline.

(43) Frost, B.; Jacks, R. L.; Diamond, M. I. Propagation of tau misfolding from the outside to the inside of a cell. J Biol Chem 2009, 284 (19), 12845–12852. DOI: 10.1074/jbc.M808759200 From NLM Medline.

(44) Khlistunova, I.; Biernat, J.; Wang, Y.; Pickhardt, M.; von Bergen, M.; Gazova, Z.; Mandelkow, E.; Mandelkow, E. M. Inducible expression of Tau repeat domain in cell models of tauopathy: aggregation is toxic to cells but can be reversed by inhibitor drugs. J Biol Chem 2006, 281 (2), 1205–1214. DOI: 10.1074/jbc.M507753200 From NLM Medline.

(45) Wille, H.; Drewes, G.; Biernat, J.; Mandelkow, E. M.; Mandelkow, E. Alzheimer-like paired helical filaments and antiparallel dimers formed from microtubule-associated protein tau in vitro. J Cell Biol 1992, 118 (3), 573–584. DOI: 10.1083/jcb.118.3.573 From NLM Medline.

(46) Grundke-Iqbal, I.; Iqbal, K.; Quinlan, M.; Tung, Y. C.; Zaidi, M. S.; Wisniewski, H. M. Microtubule-associated protein tau. A component of Alzheimer paired helical filaments. J Biol Chem 1986, 261 (13), 6084–6089. From NLM Medline.

(47) Kfoury, N.; Holmes, B. B.; Jiang, H.; Holtzman, D. M.; Diamond, M. I. Trans-cellular propagation of Tau aggregation by fibrillar species. J Biol Chem 2012, 287 (23), 19440–19451. DOI: 10.1074/jbc.M112.346072 From NLM Medline.

(48) Oakley, D. H.; Klickstein, N.; Commins, C.; Chung, M.; Dujardin, S.; Bennett, R. E.; Hyman, B. T.; Frosch, M. P. Continuous Monitoring of Tau-Induced Neurotoxicity in Patient-Derived iPSC-Neurons. J Neurosci 2021, 41 (19), 4335–4348. DOI: 10.1523/JNEUROSCI.2590-20.2021 From NLM Medline.

(49) Liu, Z.; Chen, O.; Wall, J. B. J.; Zheng, M.; Zhou, Y.; Wang, L.; Vaseghi, H. R.; Qian, L.; Liu, J. Systematic comparison of 2A peptides for cloning multi-genes in a polycistronic vector. Sci Rep 2017, 7 (1), 2193. DOI: 10.1038/s41598-017-02460-2 From NLM Medline.

(50) Zhu, X.; Ricci-Tam, C.; Hager, E. R.; Sgro, A. E. Self-cleaving peptides for expression of multiple genes in Dictyostelium discoideum. PLoS One 2023, 18 (3), e0281211. DOI: 10.1371/journal.pone.0281211 From NLM Medline.

(51) Michel, C. H.; Kumar, S.; Pinotsi, D.; Tunnacliffe, A.; St George-Hyslop, P.; Mandelkow, E.; Mandelkow, E. M.; Kaminski, C. F.; Kaminski Schierle, G. S. Extracellular monomeric tau protein is sufficient to initiate the spread of tau protein pathology. J Biol Chem 2014, 289 (2), 956–967. DOI: 10.1074/jbc.M113.515445 From NLM Medline.

(52) Gunawardana, C. G.; Mehrabian, M.; Wang, X.; Mueller, I.; Lubambo, I. B.; Jonkman, J. E.; Wang, H.; Schmitt-Ulms, G. The Human Tau Interactome: Binding to the Ribonucleoproteome, and Impaired Binding of the Proline-to-Leucine Mutant at Position 301 (P301L) to Chaperones and the Proteasome. Mol Cell Proteomics 2015, 14 (11), 3000–3014. DOI: 10.1074/mcp.M115.050724 From NLM Medline.

(53) Cherry, J. D.; Zeineddin, A.; Dammer, E. B.; Webster, J. A.; Duong, D.; Seyfried, N. T.; Levey, A. I.; Alvarez, V. E.; Huber, B. R.; Stein, T. D.;, et al. Characterization of Detergent Insoluble Proteome in Chronic Traumatic Encephalopathy. J Neuropathol Exp Neurol 2018, 77 (1), 40–49. DOI: 10.1093/jnen/nlx100 From NLM Medline.

(54) Wegmann, S.; Eftekharzadeh, B.; Tepper, K.; Zoltowska, K. M.; Bennett, R. E.; Dujardin, S.; Laskowski, P. R.; MacKenzie, D.; Kamath, T.; Commins, C.;, et al. Tau protein liquid-liquid phase separation can initiate tau aggregation. EMBO J 2018, 37 (7). DOI: 10.15252/embj.201798049 From NLM Medline.

(55) Chakraborty, P.; Riviere, G.; Liu, S.; de Opakua, A. I.; Dervisoglu, R.; Hebestreit, A.; Andreas, L. B.; Vorberg, I. M.; Zweckstetter, M. Co-factor-free aggregation of tau into seeding-competent RNA-sequestering amyloid fibrils. Nat Commun 2021, 12 (1), 4231. DOI: 10.1038/s41467-021-24362-8 From NLM Medline.

(56) Guo, Q.; Dammer, E. B.; Zhou, M.; Kundinger, S. R.; Gearing, M.; Lah, J. J.; Levey, A. I.; Shulman, J. M.; Seyfried, N. T. Targeted Quantification of Detergent-Insoluble RNA-Binding Proteins in Human Brain Reveals Stage and Disease Specific Co-aggregation in Alzheimer’s Disease. Front Mol Neurosci 2021, 14, 623659. DOI: 10.3389/fnmol.2021.623659 From NLM PubMed-not-MEDLINE.

(57) Lester, E.; Ooi, F. K.; Bakkar, N.; Ayers, J.; Woerman, A. L.; Wheeler, J.; Bowser, R.; Carlson, G. A.; Prusiner, S. B.; Parker, R. Tau aggregates are RNA-protein assemblies that mislocalize multiple nuclear speckle components. Neuron 2021, 109 (10), 1675–1691 e1679. DOI: 10.1016/j.neuron.2021.03.026 From NLM Medline.

(58) Goedert, M.; Jakes, R. Expression of separate isoforms of human tau protein: correlation with the tau pattern in brain and effects on tubulin polymerization. EMBO J 1990, 9 (13), 4225–4230. DOI: 10.1002/j.1460-2075.1990.tb07870.x.

(59) Serrano, L.; Montejo de Garcini, E.; Hernandez, M. A.; Avila, J. Localization of the tubulin binding site for tau protein. Eur J Biochem 1985, 153 (3), 595–600. DOI: 10.1111/j.1432-1033.1985.tb09342.x From NLM Medline.

(60) Nijholt, D. A.; van Haastert, E. S.; Rozemuller, A. J.; Scheper, W.; Hoozemans, J. J. The unfolded protein response is associated with early tau pathology in the hippocampus of tauopathies. J Pathol 2012, 226 (5), 693–702. DOI: 10.1002/path.3969 From NLM Medline.

(61) van der Harg, J. M.; Nolle, A.; Zwart, R.; Boerema, A. S.; van Haastert, E. S.; Strijkstra, A. M.; Hoozemans, J. J.; Scheper, W. The unfolded protein response mediates reversible tau phosphorylation induced by metabolic stress. Cell Death Dis 2014, 5 (8), e1393. DOI: 10.1038/cddis.2014.354 From NLM Medline.

(62) Ando, K.; Brion, J. P.; Stygelbout, V.; Suain, V.; Authelet, M.; Dedecker, R.; Chanut, A.; Lacor, P.; Lavaur, J.; Sazdovitch, V.;, et al. Clathrin adaptor CALM/PICALM is associated with neurofibrillary tangles and is cleaved in Alzheimer’s brains. Acta Neuropathol 2013, 125 (6), 861–878. DOI: 10.1007/s00401-013-1111-z From NLM Medline.

(63) Zhao, J.; Wu, H.; Tang, X. Q. Tau internalization: A complex step in tau propagation. Ageing Res Rev 2021, 67, 101272. DOI: 10.1016/j.arr.2021.101272 From NLM Medline.

(64) Colom-Cadena, M.; Davies, C.; Sirisi, S.; Lee, J. E.; Simzer, E. M.; Tzioras, M.; Querol-Vilaseca, M.; Sanchez-Aced, E.; Chang, Y. Y.; Holt, K.;, et al. Synaptic oligomeric tau in Alzheimer’s disease - A potential culprit in the spread of tau pathology through the brain. Neuron 2023, 111 (14), 2170–2183 e2176. DOI: 10.1016/j.neuron.2023.04.020 From NLM Medline.

(65) Kimura, T.; Whitcomb, D. J.; Jo, J.; Regan, P.; Piers, T.; Heo, S.; Brown, C.; Hashikawa, T.; Murayama, M.; Seok, H.;, et al. Microtubule-associated protein tau is essential for long-term depression in the hippocampus. Philos Trans R Soc Lond B Biol Sci 2014, 369 (1633), 20130144. DOI: 10.1098/rstb.2013.0144.

(66) Polydoro, M.; Dzhala, V. I.; Pooler, A. M.; Nicholls, S. B.; McKinney, A. P.; Sanchez, L.; Pitstick, R.; Carlson, G. A.; Staley, K. J.; Spires-Jones, T. L.;, et al. Soluble pathological tau in the entorhinal cortex leads to presynaptic deficits in an early Alzheimer’s disease model. Acta Neuropathol 2014, 127 (2), 257–270. DOI: 10.1007/s00401-013-1215-5.

(67) Chen, Q.; Zhou, Z.; Zhang, L.; Wang, Y.; Zhang, Y. W.; Zhong, M.; Xu, S. C.; Chen, C. H.; Li, L.; Yu, Z. P. Tau protein is involved in morphological plasticity in hippocampal neurons in response to BDNF. Neurochem Int 2012, 60 (3), 233–242. DOI: 10.1016/j.neuint.2011.12.013.

(68) Yin, X.; Zhao, C.; Qiu, Y.; Zhou, Z.; Bao, J.; Qian, W. Dendritic/Post-synaptic Tau and Early Pathology of Alzheimer’s Disease. Frontiers in Molecular Neuroscience 2021, 14. DOI: 10.3389/fnmol.2021.671779.

(69) Hole, K. L.; Zhu, B.; Huggon, L.; Brown, J. T.; Mason, J. M.; Williams, R. J. Tau(P301L) disengages from the proteosome core complex and neurogranin coincident with enhanced neuronal network excitability. Cell Death Dis 2024, 15 (6), 429. DOI: 10.1038/s41419-024-06815-2 From NLM Medline.

(70) Wei, Y. P.; Ye, J. W.; Wang, X.; Zhu, L. P.; Hu, Q. H.; Wang, Q.; Ke, D.; Tian, Q.; Wang, J. Z. Tau-Induced Ca(2+)/Calmodulin-Dependent Protein Kinase-IV Activation Aggravates Nuclear Tau Hyperphosphorylation. Neurosci Bull 2018, 34 (2), 261–269. DOI: 10.1007/s12264-017-0148-8 From NLM Medline.

(71) Karagoz, G. E.; Duarte, A. M.; Akoury, E.; Ippel, H.; Biernat, J.; Moran Luengo, T.; Radli, M.; Didenko, T.; Nordhues, B. A.; Veprintsev, D. B.;, et al. Hsp90-Tau complex reveals molecular basis for specificity in chaperone action. Cell 2014, 156 (5), 963–974. DOI: 10.1016/j.cell.2014.01.037 From NLM Medline.

(72) Kim, S. H.; Cho, Y. S.; Kim, Y.; Park, J.; Yoo, S. M.; Gwak, J.; Kim, Y.; Gwon, Y.; Kam, T. I.; Jung, Y. K. Endolysosomal impairment by binding of amyloid beta or MAPT/Tau to V-ATPase and rescue via the HYAL- CD44 axis in Alzheimer disease. Autophagy 2023, 19 (8), 2318–2337. DOI: 10.1080/15548627.2023.2181614 From NLM Medline.

(73) Chou, C. C.; Zhang, Y.; Umoh, M. E.; Vaughan, S. W.; Lorenzini, I.; Liu, F.; Sayegh, M.; Donlin-Asp, P. G.; Chen, Y. H.; Duong, D. M.;, et al. TDP-43 pathology disrupts nuclear pore complexes and nucleocytoplasmic transport in ALS/FTD. Nat Neurosci 2018, 21 (2), 228–239. DOI: 10.1038/s41593-017-0047-3 From NLM Medline.

(74) Toledo, J. B.; Van Deerlin, V. M.; Lee, E. B.; Suh, E.; Baek, Y.; Robinson, J. L.; Xie, S. X.; McBride, J.; Wood, E. M.; Schuck, T.;, et al. A platform for discovery: The University of Pennsylvania Integrated Neurodegenerative Disease Biobank. Alzheimers Dement 2014, 10 (4), 477–484 e471. DOI: 10.1016/j.jalz.2013.06.003 From NLM Medline.

(75) Ping, L.; Duong, D. M.; Yin, L.; Gearing, M.; Lah, J. J.; Levey, A. I.; Seyfried, N. T. Global quantitative analysis of the human brain proteome in Alzheimer’s and Parkinson’s Disease. Sci Data 2018, 5, 180036. DOI: 10.1038/sdata.2018.36 From NLM Medline.

(76) Wojtas, A. M.; Dammer, E. B.; Guo, Q.; Ping, L.; Shantaraman, A.; Duong, D. M.; Yin, L.; Fox, E. J.; Seifar, F.; Lee, E. B.;, et al. Proteomic changes in the human cerebrovasculature in Alzheimer’s disease and related tauopathies linked to peripheral biomarkers in plasma and cerebrospinal fluid. Alzheimers Dement 2024. DOI: 10.1002/alz.13821 From NLM Publisher.

(77) Seifar, F.; Fox, E. J.; Shantaraman, A.; Liu, Y.; Dammer, E. B.; Modeste, E.; Duong, D. M.; Yin, L.; Trautwig, A. N.; Guo, Q.; et al. Large-scale Deep Proteomic Analysis in Alzheimer’s Disease Brain Regions Across Race and Ethnicity. *bioRxiv* 2024. DOI: 10.1101/2024.04.22.590547 From NLM PubMed-not-MEDLINE.

(78) Braak, H.; Braak, E. Neuropathological stageing of Alzheimer-related changes. Acta Neuropathol 1991, 82 (4), 239–259. DOI: 10.1007/BF00308809 From NLM Medline.

(79) Langfelder, P.; Horvath, S. WGCNA: an R package for weighted correlation network analysis. BMC Bioinformatics 2008, 9, 559. DOI: 10.1186/1471-2105-9-559 From NLM Medline.

(80) Apicco, D. J.; Zhang, C.; Maziuk, B.; Jiang, L.; Ballance, H. I.; Boudeau, S.; Ung, C.; Li, H.; Wolozin, B. Dysregulation of RNA Splicing in Tauopathies. Cell Rep 2019, 29 (13), 4377–4388 e4374. DOI: 10.1016/j.celrep.2019.11.093 From NLM Medline.

(81) Bai, B. U1 snRNP Alteration and Neuronal Cell Cycle Reentry in Alzheimer Disease. Front Aging Neurosci 2018, 10, 75. DOI: 10.3389/fnagi.2018.00075 From NLM PubMed-not-MEDLINE.

(82) McMillan, P. J.; Strovas, T. J.; Baum, M.; Mitchell, B. K.; Eck, R. J.; Hendricks, N.; Wheeler, J. M.; Latimer, C. S.; Keene, C. D.; Kraemer, B. C. Pathological tau drives ectopic nuclear speckle scaffold protein SRRM2 accumulation in neuron cytoplasm in Alzheimer’s disease. Acta Neuropathol Commun 2021, 9 (1), 117. DOI: 10.1186/s40478-021-01219-1 From NLM Medline.

(83) Montalbano, M.; McAllen, S.; Cascio, F. L.; Sengupta, U.; Garcia, S.; Bhatt, N.; Ellsworth, A.; Heidelman, E. A.; Johnson, O. D.; Doskocil, S.;, et al. TDP-43 and Tau Oligomers in Alzheimer’s Disease, Amyotrophic Lateral Sclerosis, and Frontotemporal Dementia. Neurobiol Dis 2020, 146, 105130. DOI: 10.1016/j.nbd.2020.105130 From NLM Medline.

(84) Zaman, M.; Fu, Y.; Chen, P. C.; Sun, H.; Yang, S.; Wu, Z.; Wang, Z.; Poudel, S.; Serrano, G. E.; Beach, T. G.;, et al. Dissecting Detergent-Insoluble Proteome in Alzheimer’s Disease by TMTc-Corrected Quantitative Mass Spectrometry. Mol Cell Proteomics 2023, 22 (8), 100608. DOI: 10.1016/j.mcpro.2023.100608 From NLM Medline.

(85) Jiang, L.; Lin, W.; Zhang, C.; Ash, P. E. A.; Verma, M.; Kwan, J.; van Vliet, E.; Yang, Z.; Cruz, A. L.; Boudeau, S.;, et al. Interaction of tau with HNRNPA2B1 and N(6)-methyladenosine RNA mediates the progression of tauopathy. Mol Cell 2021, 81 (20), 4209–4227 e4212. DOI: 10.1016/j.molcel.2021.07.038 From NLM Medline.

(86) Van Nostrand, E. L.; Freese, P.; Pratt, G. A.; Wang, X.; Wei, X.; Xiao, R.; Blue, S. M.; Chen, J. Y.; Cody, N. A. L.; Dominguez, D.;, et al. A large-scale binding and functional map of human RNA-binding proteins. Nature 2020, 583 (7818), 711–719. DOI: 10.1038/s41586-020-2077-3 From NLM Medline.

(87) Bryzgalov, L. O.; Korbolina, E. E.; Merkulova, T. I. Exploring the Genetic Predisposition to Epigenetic Changes in Alzheimer’s Disease. Int J Mol Sci 2023, 24 (9). DOI: 10.3390/ijms24097955 From NLM Medline.

(88) Chiodi, V.; Domenici, M. R.; Biagini, T.; De Simone, R.; Tartaglione, A. M.; Di Rosa, M.; Lo Re, O.; Mazza, T.; Micale, V.; Vinciguerra, M. Systemic depletion of histone macroH2A1.1 boosts hippocampal synaptic plasticity and social behavior in mice. FASEB J 2021, 35 (8), e21793. DOI: 10.1096/fj.202100569R From NLM Medline.

(89) Xu, F.; Gao, W.; Zhang, M.; Zhang, F.; Sun, X.; Wu, B.; Liu, Y.; Li, X.; Li, H. Diagnostic implications of ubiquitination-related gene signatures in Alzheimer’s disease. Sci Rep 2024, 14 (1), 10728. DOI: 10.1038/s41598-024-61363-1 From NLM Medline.

(90) Wong, J. Altered expression of RNA splicing proteins in Alzheimer’s disease patients: evidence from two microarray studies. Dement Geriatr Cogn Dis Extra 2013, 3 (1), 74–85. DOI: 10.1159/000348406 From NLM PubMed-not-MEDLINE.

(91) Olah, J.; Vincze, O.; Virok, D.; Simon, D.; Bozso, Z.; Tokesi, N.; Horvath, I.; Hlavanda, E.; Kovacs, J.; Magyar, A.;, et al. Interactions of pathological hallmark proteins: tubulin polymerization promoting protein/p25, beta-amyloid, and alpha-synuclein. J Biol Chem 2011, 286 (39), 34088–34100. DOI: 10.1074/jbc.M111.243907 From NLM Medline.

(92) Nguyen, H. D.; Kim, W. K.; Huong Vu, G. Molecular mechanisms implicated in protein changes in the Alzheimer’s disease human hippocampus. Mech Ageing Dev 2024, 219, 111930. DOI: 10.1016/j.mad.2024.111930 From NLM Medline.

(93) Sengupta, U.; Montalbano, M.; McAllen, S.; Minuesa, G.; Kharas, M.; Kayed, R. Formation of Toxic Oligomeric Assemblies of RNA-binding Protein: Musashi in Alzheimer’s disease. Acta Neuropathol Commun 2018, 6 (1), 113. DOI: 10.1186/s40478-018-0615-0 From NLM Medline.

(94) Raj, T.; Li, Y. I.; Wong, G.; Humphrey, J.; Wang, M.; Ramdhani, S.; Wang, Y. C.; Ng, B.; Gupta, I.; Haroutunian, V.;, et al. Integrative transcriptome analyses of the aging brain implicate altered splicing in Alzheimer’s disease susceptibility. Nat Genet 2018, 50 (11), 1584–1592. DOI: 10.1038/s41588-018-0238-1 From NLM Medline.

(95) Diner, I.; Nguyen, T.; Seyfried, N. T. Enrichment of Detergent-insoluble Protein Aggregates from Human Postmortem Brain. J Vis Exp 2017, (128). DOI: 10.3791/55835 From NLM Medline.

(96) Hales, C. M.; Dammer, E. B.; Deng, Q.; Duong, D. M.; Gearing, M.; Troncoso, J. C.; Thambisetty, M.; Lah, J. J.; Shulman, J. M.; Levey, A. I.;, et al. Changes in the detergent-insoluble brain proteome linked to amyloid and tau in Alzheimer’s Disease progression. Proteomics 2016, 16 (23), 3042–3053. DOI: 10.1002/pmic.201600057 From NLM Medline.

(97) Bai, B.; Chen, P. C.; Hales, C. M.; Wu, Z.; Pagala, V.; High, A. A.; Levey, A. I.; Lah, J. J.; Peng, J. Integrated approaches for analyzing U1-70K cleavage in Alzheimer’s disease. J Proteome Res 2014, 13 (11), 4526–4534. DOI: 10.1021/pr5003593 From NLM Medline.

(98) Molliex, A.; Temirov, J.; Lee, J.; Coughlin, M.; Kanagaraj, A. P.; Kim, H. J.; Mittag, T.; Taylor, J. P. Phase separation by low complexity domains promotes stress granule assembly and drives pathological fibrillization. Cell 2015, 163 (1), 123–133. DOI: 10.1016/j.cell.2015.09.015 From NLM Medline.

(99) Calabretta, S.; Richard, S. Emerging Roles of Disordered Sequences in RNA-Binding Proteins. Trends Biochem Sci 2015, 40 (11), 662–672. DOI: 10.1016/j.tibs.2015.08.012 From NLM Medline.

(100) Ambadipudi, S.; Biernat, J.; Riedel, D.; Mandelkow, E.; Zweckstetter, M. Liquid-liquid phase separation of the microtubule-binding repeats of the Alzheimer-related protein Tau. Nat Commun 2017, 8 (1), 275. DOI: 10.1038/s41467-017-00480-0 From NLM Medline.

(101) Wen, J.; Hong, L.; Krainer, G.; Yao, Q. Q.; Knowles, T. P. J.; Wu, S.; Perrett, S. Conformational Expansion of Tau in Condensates Promotes Irreversible Aggregation. J Am Chem Soc 2021, 143 (33), 13056–13064. DOI: 10.1021/jacs.1c03078 From NLM Medline.

(102) Kumar, M.; Quittot, N.; Dujardin, S.; Schlaffner, C. N.; Viode, A.; Wiedmer, A.; Beerepoot, P.; Chun, J. E.; Glynn, C.; Fernandes, A. R.;, et al. Alzheimer proteopathic tau seeds are biochemically a forme fruste of mature paired helical filaments. Brain 2024, 147 (2), 637–648. DOI: 10.1093/brain/awad378 From NLM Medline.

(103) Daebel, V.; Chinnathambi, S.; Biernat, J.; Schwalbe, M.; Habenstein, B.; Loquet, A.; Akoury, E.; Tepper, K.; Muller, H.; Baldus, M.; et al. beta-Sheet core of tau paired helical filaments revealed by solid-state NMR. J Am Chem Soc 2012, 134 (34), 13982–13989. DOI: 10.1021/ja305470p From NLM Medline.

(104) von Bergen, M.; Friedhoff, P.; Biernat, J.; Heberle, J.; Mandelkow, E. M.; Mandelkow, E. Assembly of tau protein into Alzheimer paired helical filaments depends on a local sequence motif ((306)VQIVYK(311)) forming beta structure. Proc Natl Acad Sci U S A 2000, 97 (10), 5129–5134. DOI: 10.1073/pnas.97.10.5129 From NLM Medline.

(105) Younas, A.; Younas, N.; Iqbal, M. J.; Ferrer, I.; Zerr, I. Comparative interactome mapping of Tau-protein in classical and rapidly progressive Alzheimer’s disease identifies subtype-specific pathways. Neuropathol Appl Neurobiol 2024, 50 (1), e12964. DOI: 10.1111/nan.12964 From NLM Medline.

(106) Younas, N.; Zafar, S.; Saleem, T.; Fernandez Flores, L. C.; Younas, A.; Schmitz, M.; Zerr, I. Differential interactome mapping of aggregation prone/prion-like proteins under stress: novel links to stress granule biology. Cell Biosci 2023, 13 (1), 221. DOI: 10.1186/s13578-023-01164-7 From NLM PubMed-not-MEDLINE.

(107) Sinsky, J.; Majerova, P.; Kovac, A.; Kotlyar, M.; Jurisica, I.; Hanes, J. Physiological Tau Interactome in Brain and Its Link to Tauopathies. J Proteome Res 2020, 19 (6), 2429–2442. DOI: 10.1021/acs.jproteome.0c00137 From NLM Medline.

(108) Minde, D. P.; Ramakrishna, M.; Lilley, K. S. Biotin proximity tagging favours unfolded proteins and enables the study of intrinsically disordered regions. Commun Biol 2020, 3 (1), 38. DOI: 10.1038/s42003-020-0758-y From NLM Medline.

(109) Meier, S.; Bell, M.; Lyons, D. N.; Ingram, A.; Chen, J.; Gensel, J. C.; Zhu, H.; Nelson, P. T.; Abisambra, J. F. Identification of Novel Tau Interactions with Endoplasmic Reticulum Proteins in Alzheimer’s Disease Brain. J Alzheimers Dis 2015, 48 (3), 687–702. DOI: 10.3233/JAD-150298 From NLM Medline.

(110) Ayyadevara, S.; Balasubramaniam, M.; Parcon, P. A.; Barger, S. W.; Griffin, W. S.; Alla, R.; Tackett, A. J.; Mackintosh, S. G.; Petricoin, E.; Zhou, W.;, et al. Proteins that mediate protein aggregation and cytotoxicity distinguish Alzheimer’s hippocampus from normal controls. Aging Cell 2016, 15 (5), 924–939. DOI: 10.1111/acel.12501 From NLM Medline.

(111) Scheres, S. H. W.; Ryskeldi-Falcon, B.; Goedert, M. Molecular pathology of neurodegenerative diseases by cryo-EM of amyloids. Nature 2023, 621 (7980), 701–710. DOI: 10.1038/s41586-023-06437-2 From NLM Medline.

(112) Leal, C.; Zimmer, C. G. M.; Sinatti, V. V. C.; Soares, E. S.; Poppe, B.; de Wiart, A. C.; Chua, X. Y.; da Silva, R. V.; Magdesian, M. H.; Rafii, M. S.;, et al. Effects of the therapeutic correction of U1 snRNP complex on Alzheimer’s disease. Sci Rep 2024, 14 (1), 30085. DOI: 10.1038/s41598-024-81687-2 From NLM Medline.

(113) Kundinger, S. R.; Dammer, E. B.; Yin, L.; Hurst, C.; Shapley, S.; Ping, L.; Khoshnevis, S.; Ghalei, H.; Duong, D. M.; Seyfried, N. T. Phosphorylation regulates arginine-rich RNA-binding protein solubility and oligomerization. J Biol Chem 2021, 297 (5), 101306. DOI: 10.1016/j.jbc.2021.101306 From NLM Medline.

(114) Abreha, M. H.; Ojelade, S.; Dammer, E. B.; McEachin, Z. T.; Duong, D. M.; Gearing, M.; Bassell, G. J.; Lah, J. J.; Levey, A. I.; Shulman, J. M.;, et al. TBK1 interacts with tau and enhances neurodegeneration in tauopathy. J Biol Chem 2021, 296, 100760. DOI: 10.1016/j.jbc.2021.100760 From NLM Medline.

(115) Hurst, C. D.; Dunn, A. R.; Dammer, E. B.; Duong, D. M.; Shapley, S. M.; Seyfried, N. T.; Kaczorowski, C. C.; Johnson, E. C. B. Genetic background influences the 5XFAD Alzheimer’s disease mouse model brain proteome. Front Aging Neurosci 2023, 15, 1239116. DOI: 10.3389/fnagi.2023.1239116 From NLM PubMed-not- MEDLINE.

(116) Tyanova, S.; Cox, J. Perseus: A Bioinformatics Platform for Integrative Analysis of Proteomics Data in Cancer Research. Methods Mol Biol 2018, 1711, 133–148. DOI: 10.1007/978-1-4939-7493-1_7 From NLM Medline.

(117) Johnson, A. G.; Dammer, E. B.; Webster, J. A.; Duong, D. M.; Seyfried, N. T.; Hales, C. M. Proteomic networks of gray and white matter reveal tissue-specific changes in human tauopathy. Ann Clin Transl Neurol 2024. DOI: 10.1002/acn3.52134 From NLM Publisher.

(118) Dammer, E. B.; Shantaraman, A.; Ping, L.; Duong, D. M.; Gerasimov, E. S.; Ravindran, S. P.; Gudmundsdottir, V.; Frick, E. A.; Gomez, G. T.; Walker, K. A.;, et al. Proteomic analysis of Alzheimer’s disease cerebrospinal fluid reveals alterations associated with APOE epsilon4 and atomoxetine treatment. Sci Transl Med 2024, 16 (753), eadn3504. DOI: 10.1126/scitranslmed.adn3504 From NLM Medline.

(119) Guo, Q.; Ping, L.; Dammer, E. B.; Duong, D. M.; Yin, L.; Xu, K.; Shantaraman, A.; Fox, E. J.; Golde, T. E.; Johnson, E. C. B.;, et al. Heparin-enriched plasma proteome is significantly altered in Alzheimer’s disease. Mol Neurodegener 2024, 19 (1), 67. DOI: 10.1186/s13024-024-00757-1 From NLM Medline.

(120) Carey, K. M.; Young, C. D.; Clark, A. J.; Dammer, E. B.; Singh, R.; Lillard, J. W., Jr. Subtype-specific analysis of gene co-expression networks and immune cell profiling reveals high grade serous ovarian cancer subtype linkage to variable immune microenvironment. J Ovarian Res 2024, 17 (1), 240. DOI: 10.1186/s13048-024-01556-4 From NLM Medline.

(121) Saloner, R.; Staffaroni, A.; Dammer, E.; Johnson, E. C. B.; Paolillo, E.; Wise, A.; Heuer, H.; Forsberg, L.; Lago, A. L.; Webb, J.;, et al. Large-scale network analysis of the cerebrospinal fluid proteome identifies molecular signatures of frontotemporal lobar degeneration. Res Sq 2024. DOI: 10.21203/rs.3.rs-4103685/v1 From NLM PubMed-not-MEDLINE.

(122) Trautwig, A. N.; Fox, E. J.; Dammer, E. B.; Shantaraman, A.; Ping, L.; Duong, D. M.; Levey, A. I.; Lah, J. J.; Fournier, C. N.; McEachin, Z. T.;, et al. 2024. DOI: 10.1101/2024.02.29.582840.

(123) Kuchenbecker, L. A.; Thompson, K. J.; Hurst, C. D.; Opdenbosch, B. M.; Heckman, M. G.; Reddy, J. S.; Nguyen, T.; Casellas, H. L.; Sotelo, K. D.; Reddy, D. J.; et al. Nomination of a novel plasma protein biomarker panel capable of classifying Alzheimer’s disease dementia with high accuracy in an African American cohort. *bioRxiv* 2024. DOI: 10.1101/2024.07.27.605373 From NLM PubMed-not-MEDLINE.

(124) Dammer, E. B.; Ping, L.; Duong, D. M.; Modeste, E. S.; Seyfried, N. T.; Lah, J. J.; Levey, A. I.; Johnson, E. C. B. Multi-platform proteomic analysis of Alzheimer’s disease cerebrospinal fluid and plasma reveals network biomarkers associated with proteostasis and the matrisome. Alzheimers Res Ther 2022, 14 (1), 174. DOI: 10.1186/s13195-022-01113-5 From NLM Medline.

(125) Shantaraman, A.; Dammer, E. B.; Ugochukwu, O.; Duong, D. M.; Yin, L.; Carter, E. K.; Gearing, M.; Chen-Plotkin, A.; Lee, E. B.; Trojanowski, J. Q.;, et al. Network proteomics of the Lewy body dementia brain reveals presynaptic signatures distinct from Alzheimer’s disease. Mol Neurodegener 2024, 19 (1), 60. DOI: 10.1186/s13024-024-00749-1 From NLM Medline.

(126) Dammer, E. B.; Seyfried, N. T.; Johnson, E. C. B. Batch Correction and Harmonization of -Omics Datasets with a Tunable Median Polish of Ratio. Front Syst Biol 2023, *3*. DOI: 10.3389/fsysb.2023.1092341 From NLM PubMed-not-MEDLINE.

(127) Stine, R. An Introduction to Bootstrap Methods. Sociological Methods & Research 1989, 18 (2-3), 243–291. DOI: 10.1177/0049124189018002003.

(128) Seyfried, N. T.; Dammer, E. B.; Swarup, V.; Nandakumar, D.; Duong, D. M.; Yin, L.; Deng, Q.; Nguyen, T.; Hales, C. M.; Wingo, T.;, et al. A Multi-network Approach Identifies Protein-Specific Co-expression in Asymptomatic and Symptomatic Alzheimer’s Disease. Cell Syst 2017, 4 (1), 60–72 e64. DOI: 10.1016/j.cels.2016.11.006 From NLM Medline.

(129) Dai, J.; Johnson, E. C. B.; Dammer, E. B.; Duong, D. M.; Gearing, M.; Lah, J. J.; Levey, A. I.; Wingo, T. S.; Seyfried, N. T. Effects of APOE Genotype on Brain Proteomic Network and Cell Type Changes in Alzheimer’s Disease. Front Mol Neurosci 2018, 11, 454. DOI: 10.3389/fnmol.2018.00454 From NLM PubMed-not-MEDLINE.

(130) Hurst, C.; Pugh, D. A.; Abreha, M. H.; Duong, D. M.; Dammer, E. B.; Bennett, D. A.; Herskowitz, J. H.; Seyfried, N. T. Integrated Proteomics to Understand the Role of Neuritin (NRN1) as a Mediator of Cognitive Resilience to Alzheimer’s Disease. Mol Cell Proteomics 2023, 22 (5), 100542. DOI: 10.1016/j.mcpro.2023.100542 From NLM Medline.

(131) Rayaprolu, S.; Gao, T.; Xiao, H.; Ramesha, S.; Weinstock, L. D.; Shah, J.; Duong, D. M.; Dammer, E. B.; Webster, J. A., Jr.; Lah, J. J.;, et al. Flow-cytometric microglial sorting coupled with quantitative proteomics identifies moesin as a highly-abundant microglial protein with relevance to Alzheimer’s disease. Mol Neurodegener 2020, 15 (1), 28. DOI: 10.1186/s13024-020-00377-5 From NLM Medline.

(132) Levites, Y.; Dammer, E. B.; Ran, Y.; Tsering, W.; Duong, D.; Abreha, M.; Gadhavi, J.; Lolo, K.; Trejo-Lopez, J.; Phillips, J.;, et al. Integrative proteomics identifies a conserved Abeta amyloid responsome, novel plaque proteins, and pathology modifiers in Alzheimer’s disease. Cell Rep Med 2024, 5 (8), 101669. DOI: 10.1016/j.xcrm.2024.101669 From NLM Medline.

(133) Johnson, A. G.; Webster, J. A.; Hales, C. M. Glial profiling of human tauopathy brain demonstrates enrichment of astrocytic transcripts in tau-related frontotemporal degeneration. Neurobiol Aging 2022, 112, 55–73. DOI: 10.1016/j.neurobiolaging.2021.12.005 From NLM Medline.

(134) Gebauer, F.; Schwarzl, T.; Valcarcel, J.; Hentze, M. W. RNA-binding proteins in human genetic disease. Nat Rev Genet 2021, 22 (3), 185–198. DOI: 10.1038/s41576-020-00302-y From NLM Medline.

